# Bayesian inference for biophysical neuron models enables stimulus optimization for retinal neuroprosthetics

**DOI:** 10.1101/2020.01.08.898759

**Authors:** Jonathan Oesterle, Christian Behrens, Cornelius Schröder, Thoralf Herrmann, Thomas Euler, Katrin Franke, Robert G Smith, Günther Zeck, Philipp Berens

## Abstract

Multicompartment models have long been used to study the biophysical mechanisms underlying neural information processing. However, it has been challenging to infer the parameters of such models from data. Here, we build on recent advances in Bayesian simulation-based inference to estimate the parameters of detailed models of retinal neurons whose anatomical structure was based on electron microscopy data. We demonstrate how parameters of a cone, an OFF- and an ON-cone bipolar cell model can be inferred from standard two-photon glutamate imaging with simple light stimuli. The inference method starts with a prior distribution informed by literature knowledge and yields a posterior distribution over parameters highlighting parameters consistent with the data. This posterior allows determining how well parameters are constrained by the data and to what extent changes in one parameter can be compensated for by changes in another. To demonstrate the potential of such data-driven mechanistic neuron models, we created a simulation environment for external electrical stimulation of the retina as used in retinal neuroprosthetic devices. We used the framework to optimize the stimulus waveform to selectively target OFF- and ON-cone bipolar cells, a current major problem of retinal neuroprothetics. Taken together, this study demonstrates how a data-driven Bayesian simulation-based inference approach can be used to estimate parameters of complex mechanistic models with high-throughput imaging data.

## 1 Introduction

Mechanistic models have been extensively used to study the biophysics underlying information processing in single neurons and small networks in great detail^1, 2^. In contrast to phenomenological models used for neural system identification, such models try to preserve certain physical properties of the studied system to facilitate interpretation and a causal understanding. For example, biophysical models can incorporate the detailed anatomy of a neuron^3^, its ion channel types^4, 5^ and the distributions of these channels^6^ as well as synaptic connections to other cells^7^. For all these properties, the degree of realism can be adjusted as needed. While increased realism may enable models to capture the highly non-linear dynamics of neural activity more effectively, it usually also increases the number of model parameters. While the classical Hodgkin-Huxley model with one compartment has already ten free parameters^4^, detailed multicompartment models of neurons can have dozens or even hundreds of parameters^8^.

Constraining many of these model parameters such as channel densities requires highly specialized and technically challenging experiments, and, hence, it is usually not viable to measure every single parameter for a neuron model of a specific neuron type. Rather, parameters for mechanistic simulations are often aggregated over different neuron types and even across species. Even though this may be justified in specific cases it likely limits our ability to identify mechanistic models of individual cell types. Alternatively, parameter search methods have been proposed to identify the parameters of mechanistic neuron models from standardized patch-clamp protocols based on exhaustive grid-searches^9–11^ or evolutionary algorithms^12–15^. Such methods are often inefficient, typically not applicable for models with many parameters and identify only a single point estimate consistent with the data instead of the entire distribution.

Here, we built on recent advances in Bayesian simulation-based inference to fit multicompartment models of neurons with realistic anatomy in the mouse retina. We used a framework called Sequential Neural Posterior Estimation (SNPE)^16, 17^ to identify model parameters based on high-throughput two-photon measurements of these neurons’ responses to light stimuli. SNPE is a Bayesian simulation-based inference algorithm that allows parameter estimation for simulator models for which the likelihood cannot be evaluated easily. The algorithm estimates the distribution of model parameters consistent with specified target data by evaluating the model for different sets of parameters and comparing the model output to the target data. To this end, parameters are drawn from a prior distribution, which is an initial guess about which parameters are likely to produce the desired model output. For example, the choice of prior distribution can be informed by the literature, without constraining the model to specific values. The model output for the sampled parameter sets can than be used to refine the distribution over plausible parameters given the data. This updated distribution, containing information from both the prior and the observed simulations, is known as the posterior. For high dimensional parameter spaces, many samples are necessary to obtain an informative posterior estimate. Therefore, to make efficient use of simulation time, SNPE iteratively updates its sampling distribution, such that only in the first round samples are drawn from the prior, while in subsequent rounds samples are drawn from intermediate posteriors. This procedure increases the fraction of samples leading to simulations close to the target data. Since this approach for parameter estimation not only returns a point-estimate but also a posterior distribution over parameters consistent with the data, it allows one to straightforwardly determine how well the parameters are constrained. While the method has been used previously to fit simple neuron models^16, 17^, it has so far not been applied to models as complex and realistic as the ones presented here.

We estimated the posterior parameter distribution of multicompartment models of three retinal neurons, a cone photoreceptor (cone), an OFF- and an ON-bipolar cell (BC). The structure of the BC models was based on high-resolution electron microscopy reconstructions^18^ and in eight independently parameterized regions. We performed parameter inference based on the responses of these neurons to standard light stimuli measured with two-photon imaging of glutamate release using iGluSnFR as an indicator^19^. Our analysis shows that many of the model parameters can be constrained well, yielding simulation results consistent with the observed data. After validating our model, we show that the inferred models and the inference algorithm can be used to efficiently guide the design of electrical stimuli for retinal neuroprosthetics to selectively activate OFF- or ON-BCs. This is an important step towards solving a long-standing question in the quest to provide efficient neuroprosthetic devices for the blind.

## 2 Methods

### 2.1 Biophysical neuron models

We created detailed models of three retinal cell types: a cone, an ON- (Fig. 1A, Bi) and an OFF-BC (Fig. 1Bii). From the different OFF- and ON-BC types we chose to model the types 3a and 5o, respectively, because those were the retinal cone bipolar cell (CBC) types in mice for which we could gather most information. To model the light response, the OFF-BC model received input from five and the ON-BC from three cones^20^. Every cone made two synaptic connection with each BC.

**Figure 1.**
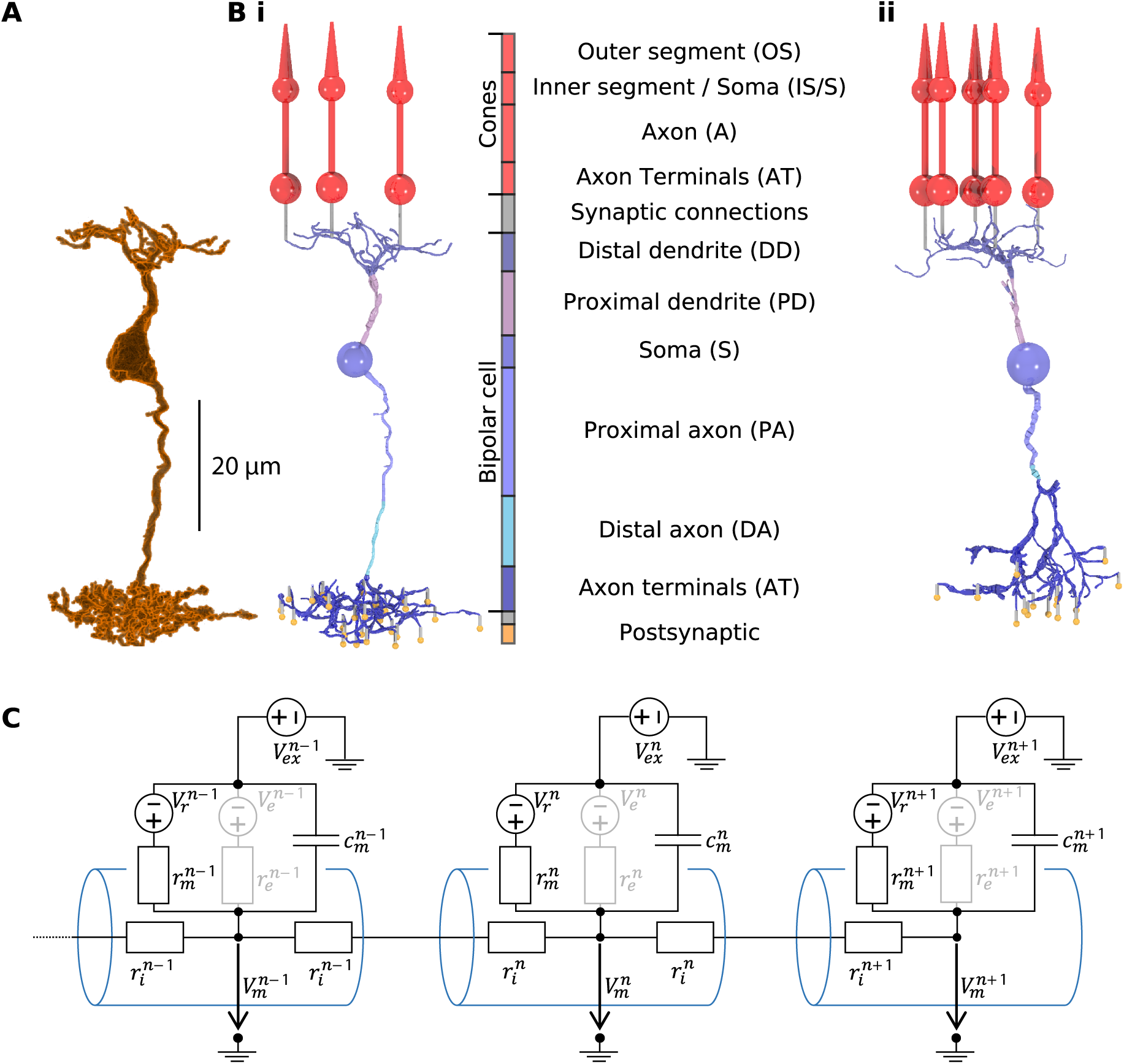
From serial block-face electron microscopy (EM) data of retinal BCs to multicompartment models. **(A)** Raw morphology extracted from EM data of an ON-BC of type 5o. **(Bi)** Processed morphology connected to three presynaptic cones (red) and several postsynaptic compartments (yellow). The cone and BC morphologies are divided into color-coded regions with a legend shown on the right. **(Bii)** Same as (Bi) but for an OFF-BC of type 3. **(C)** Three cylindrical compartments of a multicompartment model. Every compartment (blue) n consists of a membrane capacitance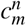, a membrane resistance 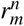, a leak conductance voltage source 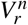, an extracellular voltage source 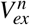 and at least one axial resistor 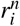 that is connected to a neighboring compartment. 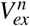 is only used to simulate electrical stimulation and is otherwise replaced by a shortcut. Compartments may have one or more further voltage- or ligand-dependent resistances 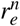 with respective voltage sources 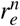 to simulate ion channels (indicated in grey).

#### Multicompartment models

We used NeuronC^21^ to implement multicompartment models of these neurons. A multicompartment model subdivides a neuron into a finite number of compartments. Every compartment is modeled as an electrical circuit, has a position in space, a spatial shape and is connected to at least one neighboring compartment (Fig. 1C). The voltage in a compartment *n*, connected to the compartments *n* − 1 and *n* + 1 is described by:

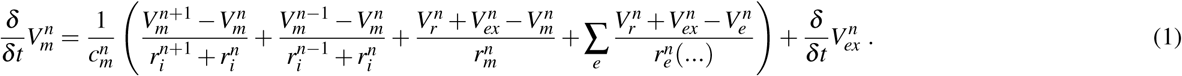

Here, compartments are either modeled as cylinders or spheres. The membrane capacitance *c*_*m*_, membrane resistance *r*_*m*_ and axial resistance *r*_*i*_ are assumed to be dependent on the compartment surface area *A*_*m*_ and/or the compartment length *l*_*c*_. The respective values for a compartment *n* are computed as:

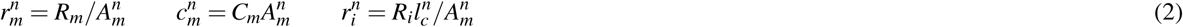

We assumed the area specific membrane resistance *R*_*m*_, the area specific membrane capacitance *C*_*m*_ and the axial resistivity *R*_*i*_ to be constant over all compartments within a cell model. We used *R*_*i*_ = 132Ωcm for all cell types and informed our priors for *C*_*m*_ and *R*_*m*_, which we estimated for every cell type individually, based on measurements from rod bipolar cells of rats^22^.

#### Anatomy

We used a simplified cone morphology consisting of four compartments: one cone shaped compartment for the outer segment, one spherical compartment for the combination of inner segment and soma, one cylindrical compartment for the axon and another spherical one for the axonal terminals (Fig. 1). The light collecting area in the outer segment was set to 0.2 µm^2^ ^23^. The diameter of the soma 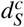, the axon 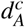 and axonal terminals 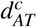, the length of the axon 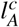 and the length of the outer segment 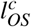 were based on electron microscopy data ^24^:

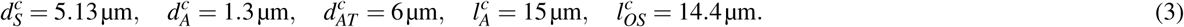

The BC morphologies in this study were based on serial block-face electron microscopy data of mouse bipolar cells^18^. We extracted the raw voxel-based morphologies from the segmentation of the EM dataset and transformed them into a skeleton plus diameter representation using Vaa3D-Neuron2 auto tracing^25^. These where then manually refined using Neuromantic^26^ to correct errors originating from small segmentation errors (Fig. 1). The ON-BC morphology we chose was classified as type 5o, equal to the functional type of the model. For the OFF-BC we decided for a morphology classified as type 3b although we functionally modeled a type 3a cell, because the chosen reconstructed morphology was of higher quality than all available type 3a reconstructions and because type 3a and 3b BCs have very similar morphologies. Additionally, type 3a and 3b mostly differ in the average axonal field size^18, 27^, with that of type 3a being larger than that of type 3b, and the selected morphology has the largest axonal field among all cells classified as 3b in the dataset, well within the range of type 3a cells.

Because the computational time scales approximately linear with the number of BC compartments, using the full number of compartments of the EM reconstructions (> 1000) during parameter inference was computationally infeasible. Therefore, we utilized the compartment condensation algorithm of NeuronC, which iteratively reduces the number of compartments while preserving biophysical properties^21^. To be able to draw a sufficient number of samples, we reduced the number of compartments during parameter inference to 10 and 13 for the OFF- and ON-BC respectively (requiring ≈4 min per simulation for a 30 s light stimulus). To simulate the electrical stimulation, more compartments are necessary to capture the effect of the electrical field on the neurites of the BC models. Therefore, we increased the number of compartments to 97 and 101 for the OFF- and an ON-BC, respectively, which is sufficient to accurately represent all major neurites without becoming computationally too expensive (requiring ≈65 and 26 min per simulation for a 30 s light stimulus for the OFF- and ON-BC respectively).

#### Ion channels and synapses - Literature review

The complement and distribution of voltage- or ligand-gated ion channels shapes the response of neurons. Here, ion channels are modeled as additional electrical elements in the compartments’ membrane with conductances dependent on time varying parameters, such as the membrane potential and the calcium concentration within the cell. In addition to the equations that govern a channel’s kinetics, their location in the cell has to be defined. After a literature review of retinal cone bipolar cell types in mice, we decided to model the OFF- and ON-type for which we could gather most information, namely BC3a and BC5. Currently, there are three accepted subtypes of BC5, namely 5o, 5i and 5t^28^. Here, we modeled the BC5 subtype that expresses voltage-gated sodium channels^29^ which probably also corresponds to the more transient BC5 subtype reported in ^30^. The TTX sensitivity observed in ^31^ suggests that both, 5o and 5i express voltage-gated sodium channels. To make our model consistent, we used data from the same BC5 subtype (5o) for the morphology, the target data and the number of cone contacts. A summary of all used channels, their location within the models and the respective references can be found in Table 1. The following paragraphs describe which channels were included in the models and why. Note however, that for all channels (except the L-type calcium channel in the axon terminals, as calcium channels are necessary in the model for neurotransmitter release) channel densities of zero were included in the prior distributions, thereby allowing the parameter inference to effectively remove ion channels from the model.

**Table 1.** Ion channels of biophysical models.

| Channel | Cone | OFF-BC (type 3a) | ON-BC (type 5) | Cone references | BC references |
| --- | --- | --- | --- | --- | --- |
| Ca<sub>L</sub> | AT | S, AT | S, AT | 32,33 | 36 |
| Ca<sub>T</sub> |  | S, AT |  |  | 36 |
| Ca<sub>p</sub> | AT | S, AT | S, AT | 34 | 34 |
| HCN<sub>1</sub> | All |  | D, S, AT | 35,36 | 29,35 |
| HCN<sub>4</sub> |  | D, S, AT |  |  | 29,35 |
| K<sub>v</sub> | IS/S | DD, PA, DA | DD, PA, DA | 35,36 | 42 |
| K<sub>ir</sub> |  | S | S |  | 35,40 |
| Cl<sub>Ca</sub> | AT |  |  | 38,39 |  |
| Na<sub>v</sub> |  | DA | DA |  | 29 |

Regions of ion channels and the respective abbreviations as in Fig. 1.
D refers to the combination of DD and PD. All refers to the combination of IS/S, A and AT.
If multiple regions are stated for a single neuron, the ion channel density differs between them.

In their axon terminals, cones express L-type calcium (Ca_L_) channels that mediate release of the transmitter glutamate^32, 33^. We modeled calcium extrusion purely with calcium pumps (Ca_P_) since other mechanisms such as sodium-calcium-exchangers probably only play a minor functional role in cones^34^. Additionally, there is evidence that cones express hyperpolarization-activated cyclic nucleotide-gated cation (HCN) channels of the type 1, mostly in the inner segment but also in the axon^35, 36^. The presence of HCN_3_ channels in mouse cones is more controversial. These channels have been observed in rat cones^37^, and a more recent study also found evidence for HCN_3_ channels at the synaptic terminals of mouse cones, but could not observe any functional differences between wild-type and HCN_3_ -knockout mice. To restrict the number of model parameters, we did not include HCN_3_ in our cone model. However, we added calcium-activated chloride (Cl_Ca_) channels to the axon terminals, based on findings in ^38, 39^ and voltage-gated potassium channels K_*V*_ at the inner segment^36^.

Our BC5 type expresses voltage-gated sodium (Na_V_) channels at the axon shaft^29^. Another study found inward-rectifier potassium (K_ir_) channels at the soma of BC5^35^, that were also found in the homologous type in rat^40^. Additionally, BC5 express HCN channels at the axon terminal, the soma and the dendrites^29, 35^. From the four subtypes of HCN, BC5 seem to almost exclusively express HCN_1_. In the rat, there is also evidence for the expression of HCN_4_ channels in BC5^37, 41^, but this could not be verified for mice. Data from rat suggests that BCs with Na_V_ channels also express K_V_ channels^42^. We therefore added K_V_ channels at the dendrites and the axon.

Similar to BC5, BC3a express HCN channels at the axon terminals, the soma and the dendrites. However, instead of HCN_1_ they express HCN_4_^29, 35^. There is also evidence that BC3a express Na_V_ channels at the axon shaft^29^ which were also found in the homologous type in rat^40^. Just like for BC5 we added also K_V_. K_ir_ in BC3a were only reported for rat so far^40^. Since we could not find any evidence for the lack of K_ir_ channels in mouse BC3a and the channel repertoires of BC3a in mouse and rat are overall very consistent, we included them in our model.

The distribution of calcium channels in mouse CBCs is largely unknown^36^. In the rat retina, there is evidence for T-type calcium (Ca_T_) channels in BC3a^41^. Calcium currents of unspecified type were observed in BC5^40^. Generally, L-type calcium (Ca_L_) channels are believed to mediate neurotransmitter release in almost all BCs across types and species^36^. Therefore, we included them in both BC models. The literature review in ^36^ suggests that T-type calcium channels might be exclusively expressed in BC3. In mouse BC3b, the simultaneous expression of both Ca_T_ and Ca_L_ has been described^43^. Furthermore, the latter and other studies^44, 45^ suggest that voltage-gated calcium channels might not be located in the axon terminals only, but also in the soma and might play a role in signal transmission within the cell. Based on the studies mentioned, we assumed that BC3a and BC5 express Ca_L_ in the axon terminals and potentially also at the soma. The BC3a model may additionally use Ca_T_ channels, both at the soma and at the axon terminals. For calcium extrusion, we added calcium pumps^34^.

BC5 receive input from cones via the metabotropic glutamate receptor 6 (mGluR6)^36^. BC3a receive input from cones via kainate receptors^46^. We modeled the kainate receptors by modifying the inactivation time constant *τ*_*γ*_ of the AMPA receptors included with NeuronC.

#### Ion channels and synapses - Implementation

All ion channels in this study were based on the models available in NeuronC. We used both Hodgkin-Huxley (HH) and Markov-Sequential-States (MS) channel implementations. Since we did not add channel noise to our model, every HH channel could have also been described as an equivalent MS channel. However, since HH channels are computationally less expensive, we used HH implementations wherever possible. Implementation details and references are listed in Table 2. The L-type calcium channel, for example, was based on the HH model defined by the following equations:

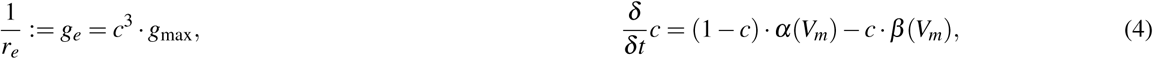

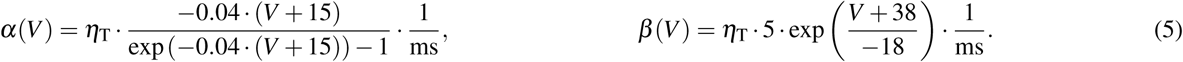

**Table 2.**
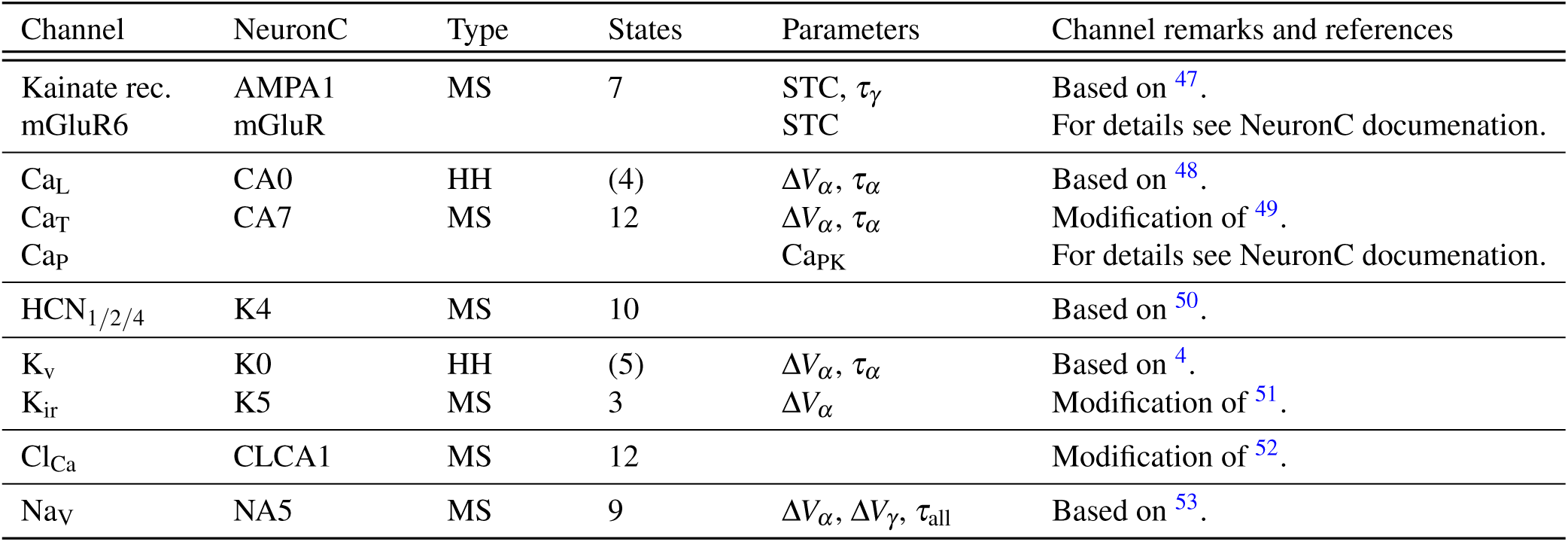
Ion channel implementation details and optimized channel parameters.

**Table 3.** Parameters of discrepancy measures.

| Discrepancy | Cone |  |  |  | BC (3a \| 5o) |  |  |  | References |
| --- | --- | --- | --- | --- | --- | --- | --- | --- | --- |
| | $p_l$ | $t_l$ | $t_u$ | $p_u$ | $p_l$ | $t_l$ | $t_u$ | $p_u$ | |
| $\delta_{\text{Rate}}^{\text{Rest}}$ | 0 | 50 | 80 | 100 | 0 | 0 | 3 \| 4 | 7 | 57,60,67,68 |
| $\delta_{V_m}^{\text{Rest}}$ | -80 | -55 | -40 | -20 | -80 | -65 | -45 | -30 | 30,42,46,66 |
| $\delta_{\text{Rate}}^{\Delta}$ | 0 | 50 | 65 | 100 | 0 | 5 | ∞ | ∞ | |
| $\delta_{V_m}^{\Delta}$ | 0 | 5 | 10 | 20 | 0 | 5 | 15 \| 25 | 40 | 30,46 |
| $\delta_{V_m}^{\text{min}}$ | -75 | 60 | ∞ | ∞ | -100 | -80 | ∞ | ∞ | 42,66,69 |
| $\delta_{V_m}^{\text{max}}$ | $-\infty$ | $-\infty$ | -35 | -20 | $-\infty$ | $-\infty$ | -10 | 0 | 42,66,69 |

Here, *η*_T_ corrects for differences between the temperature of the simulated cell *T*_sim_ and the temperature for which the channel equations were defined *T*_eq_ based on a temperature sensitivity *Q*_10_ which can vary between ion channels and state transitions:

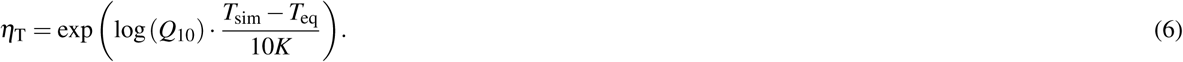

There are several sources for model uncertainty about the exact channel kinetics. First, not all channel models used here were developed based on mouse data resulting in species dependent differences. Second, we do not always know the exact subtypes of ion channels, e.g. in the case of the T-type calcium channel. Third, the exact temperature sensitivities *Q*_10_ are not known. Therefore, we estimated transition rates and thresholds for state transitions during the parameter inference. For this, we allowed for offsets Δ*V* relative to *V*_*m*_ in the rate equations and additionally, we estimated relative time constants *τ* for the rates. For example 4 was changed to:

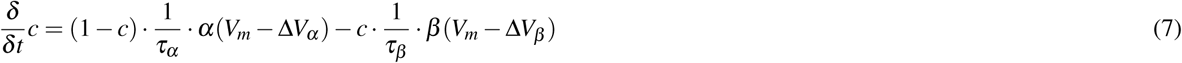

To keep the parameter space as small as possible, we only optimized the kinetics of ion channels with high uncertainty (e.g. K_v_) or with high relevance for the exact timing of the neurotransmitter release (e.g. Ca_L_ and Ca_T_). Additionally, we constrained the channel parameters to physiologically plausible ranges. Table 2 summarizes which channel parameters were estimated during parameter optimization. Time constants *τ* and voltage offsets Δ*V* not optimized were set to one and zero, respectively. For the Na_V_, a single time constant *τ*_all_ was used to modify all time constants proportionally. The calcium pump dynamics were modified by changing the calcium concentration Ca_PK_ that causes half of the maximum calcium extrusion velocity. The BC glutamate receptors were optimized by allowing for a change in the synaptic transmitter concentration at the receptors by a factor of STC, which might be smaller for the OFF-BC than for the ON-BC given the greater distance between the release sites of the cones and the dendritic tips of the BCs^20^. The simulated cell temperature *T*_sim_ was set to 37 °C if not stated otherwise. For further information we refer to the NeuronC documentation^21^.

#### Neurotransmitter release

Both cones and BCs release glutamate from ribbon synapses in response to calcium influx^54, 55^. We modeled the ribbon synapses with a standard model^21^ including a readily releasable pool (RRP) from which vesicles can be released^56^. The release rate is dependent on the number of vesicles currently available *v*_*RRP*_ in the RRP, the maximum number of vesicles 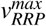 in the RRP and the intracellular calcium concentration [*Ca*]. In NeuronC, calcium is modeled in radial shells through which calcium can diffuse deeper into the neuron. For the release of neurotransmitter, only the calcium concentration in the first shell [*Ca*]_0_ (equivalent to the concentration at the membrane) is considered. The release rate *r* is computed as:

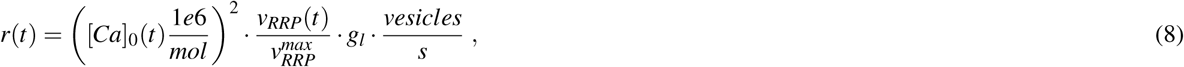

where *g*_*l*_ is a linear gain factor. *g*_*l*_ and 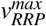 were optimized for every cell type individually. The RRP is constantly replenished with a constant rate that is equivalent to the maximum sustainable release rate *r*_*msr*_. At a time *t*, for a simulation time step Δ*t*, the vesicles in the pool are updated as follows:

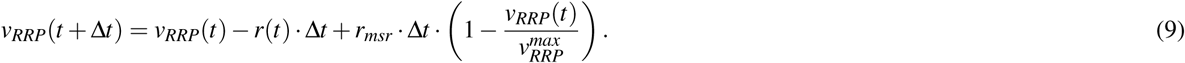

For the cone model, *r*_*msr*_ was set to 100 vesicles per second based on ^57^. The prior for 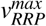 was based on RRP sizes reported for salamander^58, 59^. For the BCs, *r*_*msr*_ was set to 8 vesicles per second based on the reported value for rat rod bipolar cells in^60^. The prior for 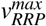 was based on ^61^.

### 2.2 Bayesian inference for model parameters

To estimate the free parameters of the multicompartment models, we used a Bayesian likelihood-free inference framework called Sequential Neural Posterior Estimation (SNPE). The goal of the parameter estimation was to find model parameters which yield model outputs matching the experimentally observed glutamate release in response to a light stimulus (Fig. 5A and 6A). Details of the algorithm, the target data, the stimulus and the comparison between experimental and simulated data are described below. To be able to simulate the light response of the BC models, we estimated the parameters of the cone model first.

**Figure 2.**
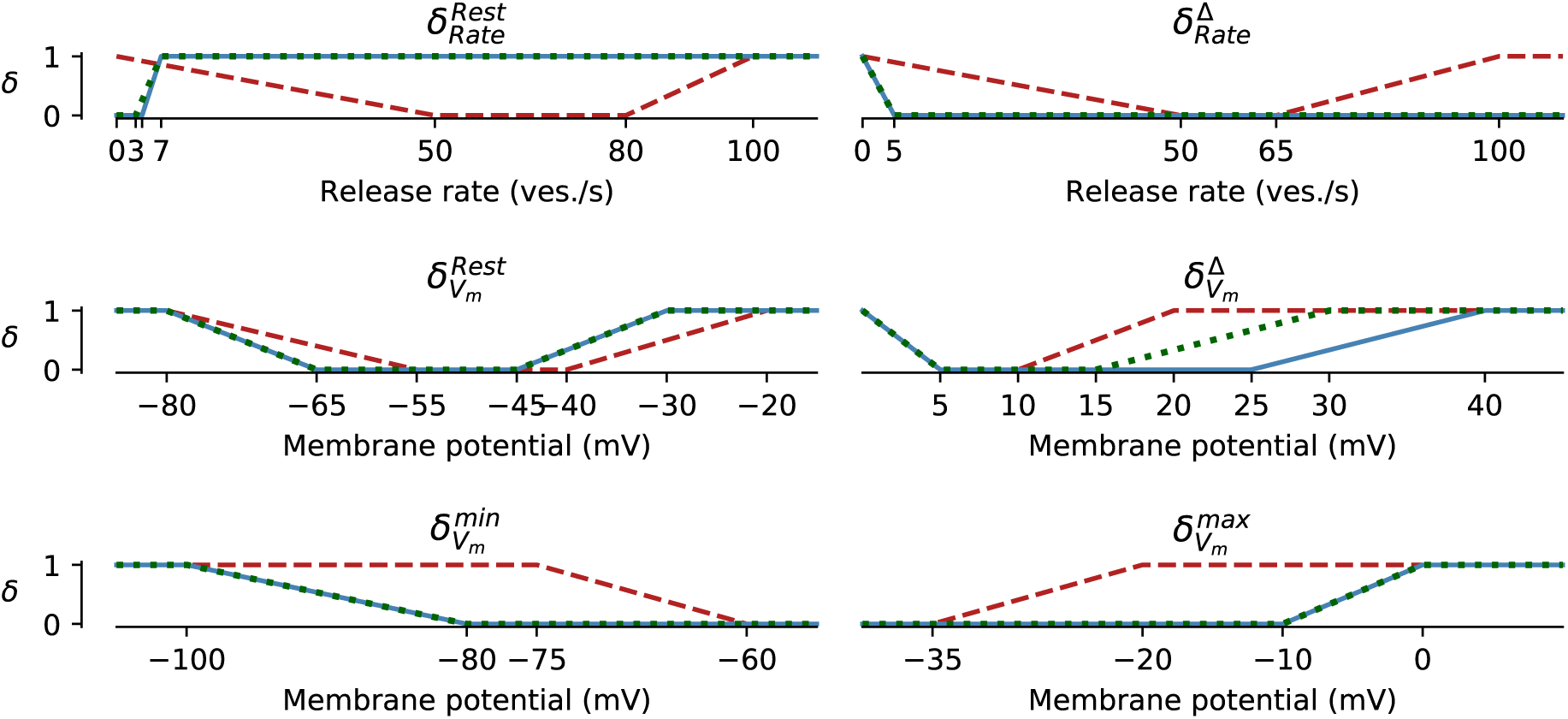
Discrepancy measures based on equation 15 for the cone (red dashed line), the OFF- (blue solid line) and ON-BC (green dotted line). The parameters defining the discrepancy measures are listed Table 3. All discrepancy measures are between zero and one per definition.

**Figure 3.**
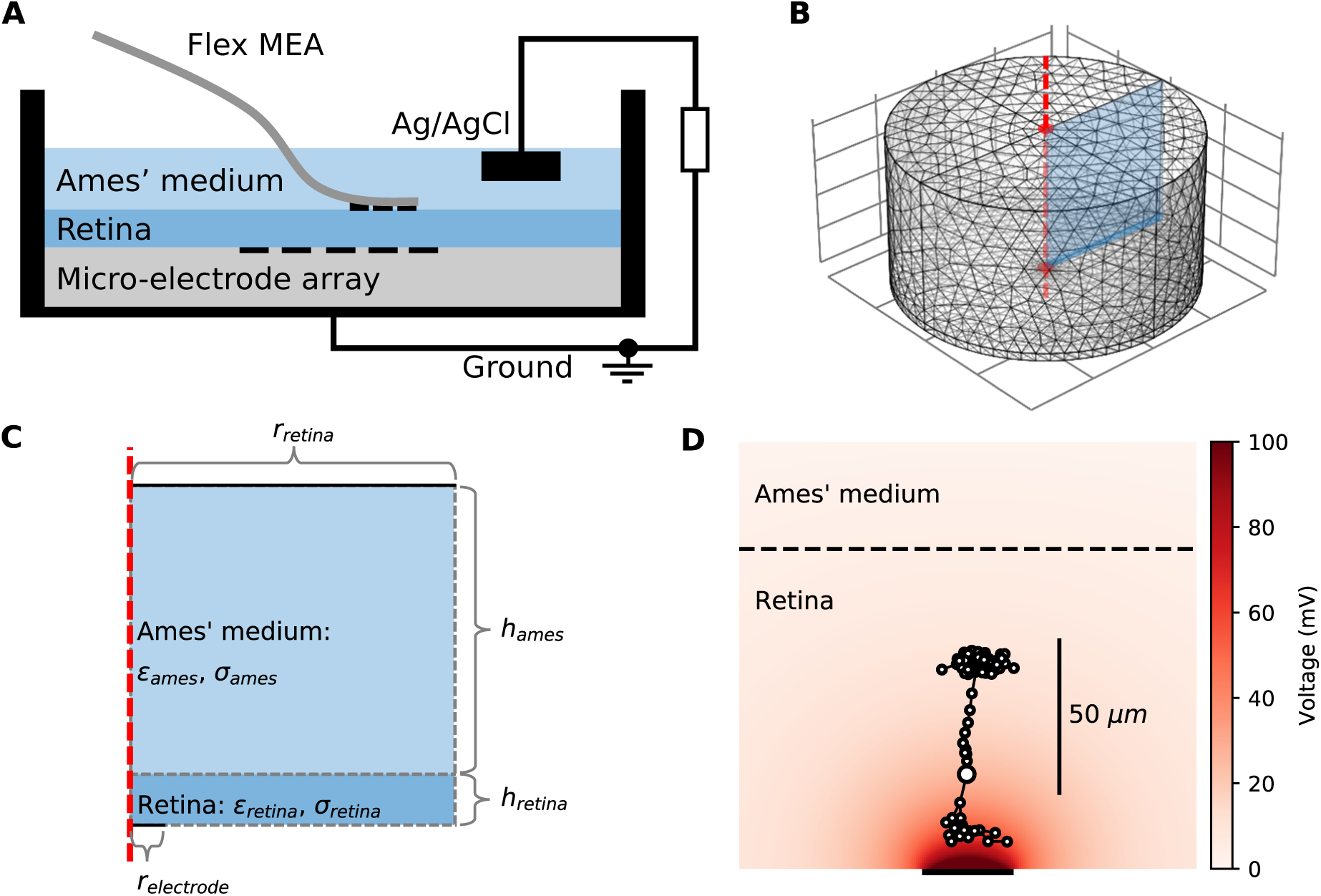
*Model for the external electrical stimulation of the retina*. ***(A)*** *Schematic figure of the experimental setup for subretinal stimulation of ex vivo retina combined with epiretinal recording of retinal ganglion cells. Schematic modified from* ^*72*^. ***(C, D)*** *Model for simulating the electrical field potential in the retina in 3D and 2D, respectively. The retina (darker blue) and the electrolyte above (lighter blue) are modeled as cylinders. The shown 3D model is radially symmetric with respect to the central axis (red dashed line). Therefore, the 3D and 2D implementations are equivalent, except that the computational costs for the 2D model are much lower. The 2D implementation is annotated with parameters that were either taken from the literature or inferred from experimental data*. ***(D)*** *Electrical field potential in the retina for a constant stimulation current of* 0.5 µA *for a single stimulation electrode with a radius of* 15 µm. *Additionally, the compartments (black circles with white filling) of the ON-BC model are shown. The stimulation is subretinal meaning that the dendrites are facing the electrode (horizontal black line on bottom)*.

**Figure 4.**
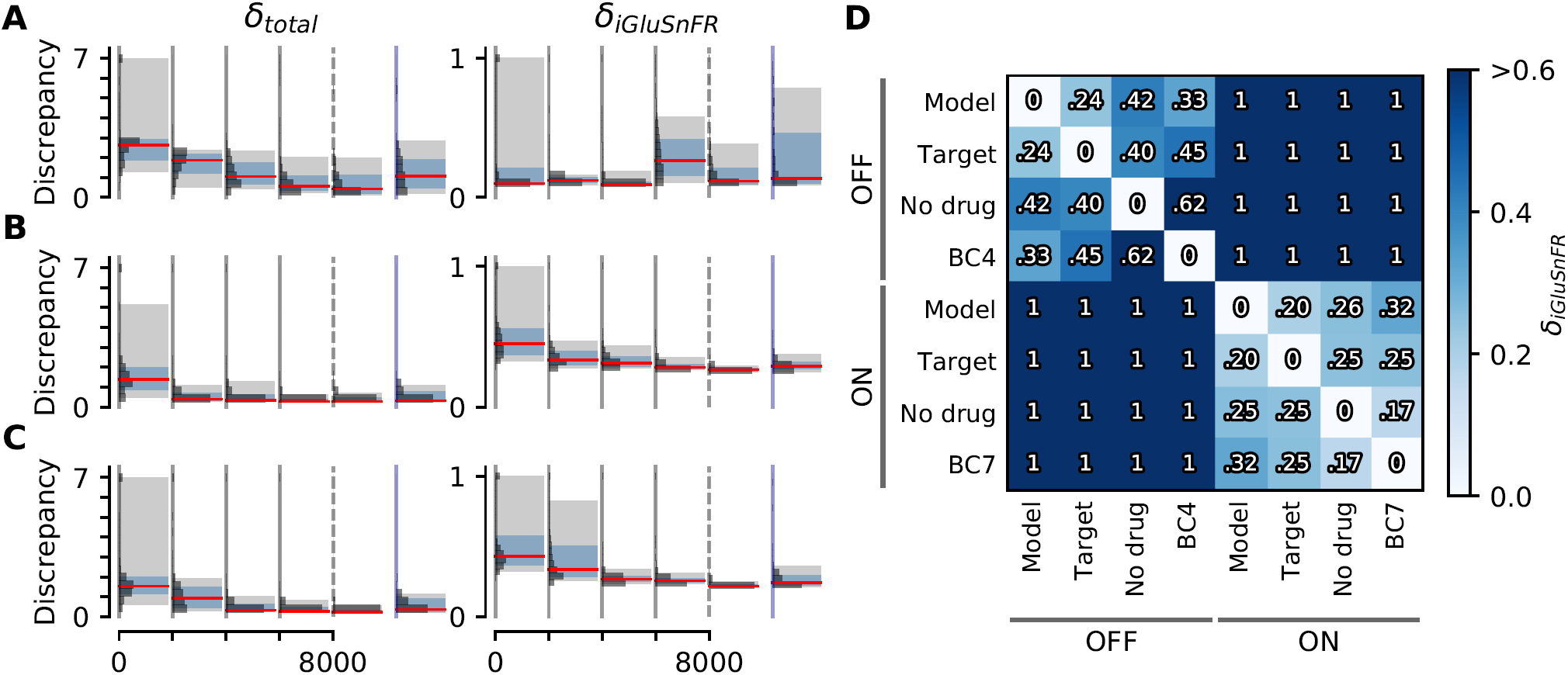
*Discrepancies of samples from the cone and the BC models during and after parameter estimation*. ***(A, B, C)*** *Sampled discrepancies for the cone (top), the OFF- (center), and ON-BC (bottom) respectively. For every model, the total discrepancy δ*_*tot*_ *(left) and the discrepancy between the simulated and target iGluSnFR trace δ*_*iGluSnFR*_ *(right) are shown. For every model, four optimization rounds with* 2, 000 *samples each were drawn (indicated by gray vertical lines). After the last round (indicated by dashed vertical lines)*, 200 *more samples were drawn from the posteriors. For the BCs, the number of compartments was increased in this last step to* 97 *and* 101 *for the OFF- and ON-BC respectively. Additionally*, 200 *samples were drawn from assuming independent posterior marginals for comparison (indicated by blue vertical lines). For every round, the discrepancy distribution (horizontal histograms), the median discrepancies (red vertical lines), the 25th to 75th percentile (blue shaded area) and the 5th to 95th percentile (gray shaded area) are shown*. ***(D)*** *Discrepancies between different iGluSnFR traces of BCs to demonstrate the high precision of the model fit. The pairwise discrepancy computed with equation 14 between eight iGluSnFR traces is depicted in a heat map. The column and row labels indicate which* ***x***_*t*_ *and* ***x***_*s*_ *were used in equation 14 respectively. The traces consists of the optimized BC models (“Model”), the targets used during optimization (“Target”), experimental data from the same cell type without the application of any drug (“No drug”) and experimental data from another retinal CBC type with the application of strychnine (“BC4” and “BC7”). Note that strychnine was also applied to record the targets*.

**Figure 5.**
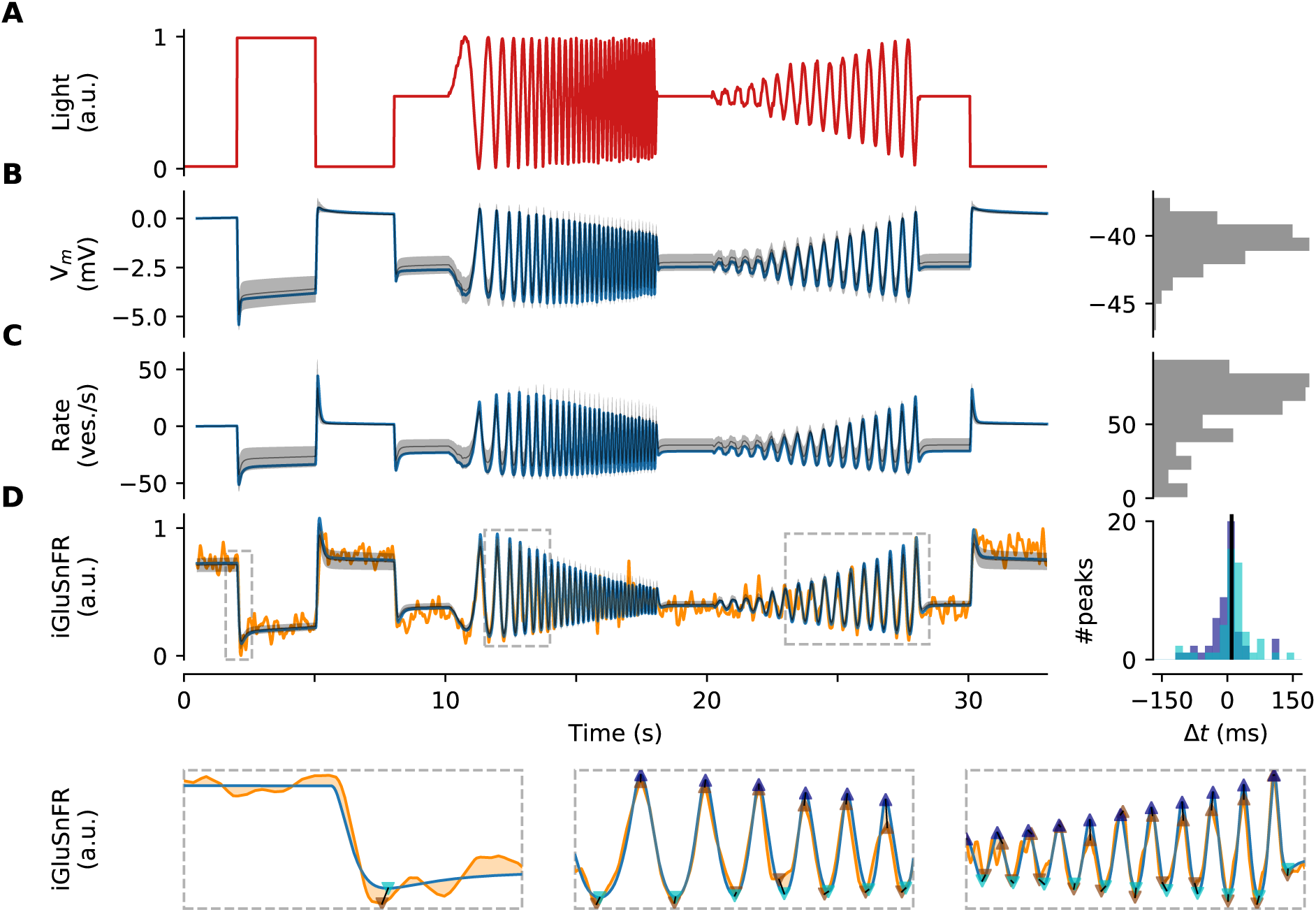
*Optimized cone model*. ***(A)*** *Normalized light stimulus*. ***(B)*** *Somatic membrane potential relative to the resting potential for the best parameters (blue line) and for* 200 *samples from the posterior shown as the mean (gray dashed line) plus-minus one standard-deviation (gray shaded area). A histogram over all resting potentials is shown on the right*. ***(C)*** *Release rate relative to the resting rate. Otherwise as in (B)*. ***(D)*** *Simulated iGluSnFR trace (as in (B)) compared to target trace (orange). Three regions (indicated by gray dashed boxes) are shown in more detail below without samples from the posterior. Estimates of positive and negative peaks are highlighted (up- and downward facing triangles respectively) in the target (brown) and in the simulated trace (blue and cyan respectively). Pairwise time differences between target and simulated peaks (indicated by triangle pairs connected by a black line) are shown as histograms for positive (blue) and negative (cyan) peaks on the right. The median over all peak time differences is shown as a black vertical line*.

**Figure 6.**
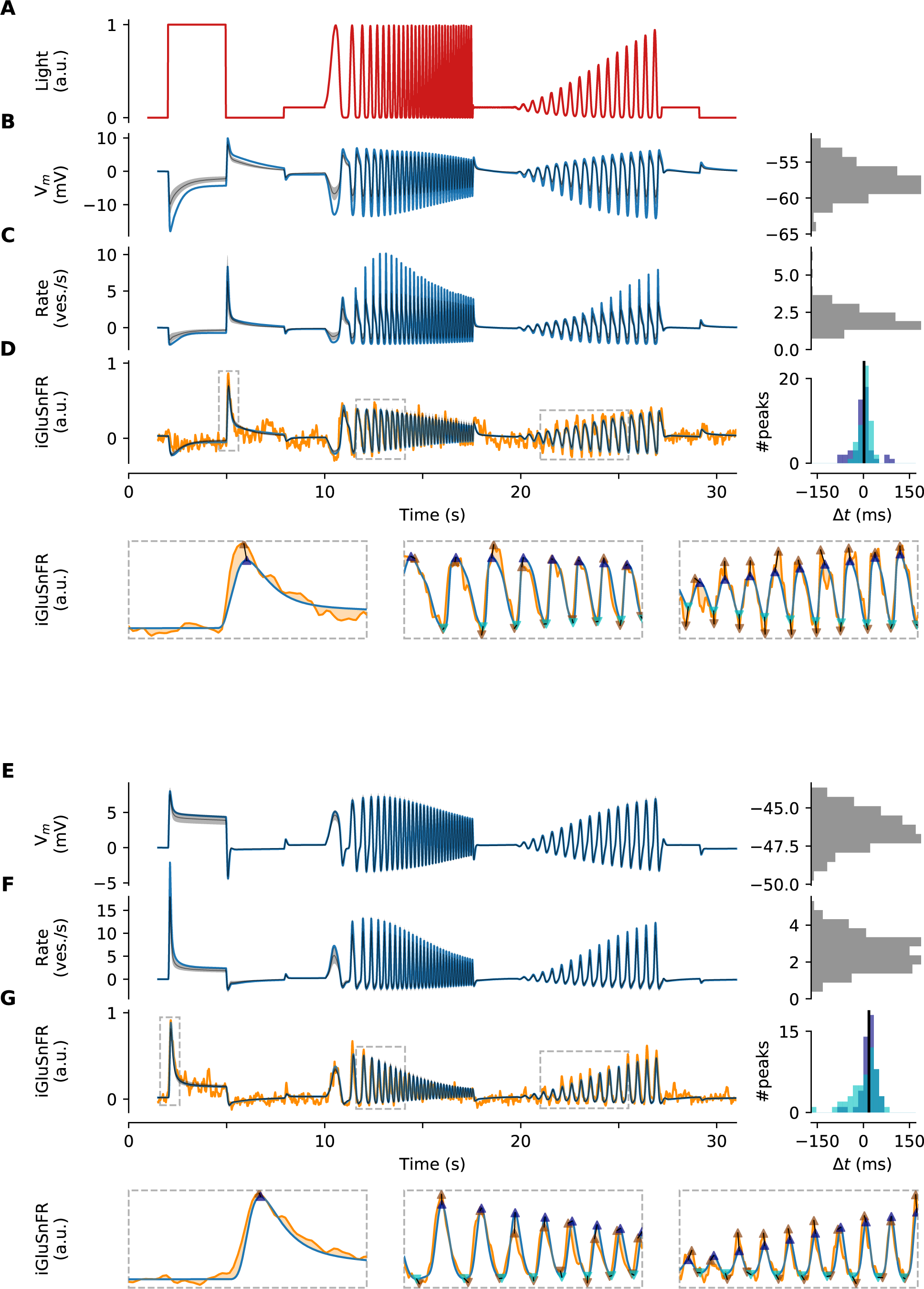
*Optimized BC models*. ***(A)*** *Normalized light stimulus. Responses of the OFF- and ON-BC are shown in (B-D) and (E-G), respectively*. ***(B, E)*** *Somatic membrane potential relative to the resting potential for the best parameters (blue line) and for* 200 *samples from the posterior shown as the mean (gray dashed line) plus-minus one standard-deviation (gray shaded area). A histogram over all resting potentials is shown on the right*. ***(C, F)*** *Mean release rate over all synapses relative to the mean resting rate. Otherwise as in (B)*. ***(D, G)*** *Simulated iGluSnFR trace (as in (B)) compared to respective target trace (orange). Three regions (indicated by gray dashed boxes) are shown in more detail below without samples from the posterior. Estimates of positive and negative peaks are highlighted (up- and downward facing triangles respectively) in the target (brown) and in the simulated trace (blue and cyan respectively). Pairwise time differences between target and simulated peaks (indicated by triangle pairs connected by a black line) are shown as histograms for positive (blue) and negative (cyan) peaks on the right. The median over all peak time differences is shown as a black vertical line*.

#### Priors

As every Bayesian method, the inference algorithm needs a prior distribution *p*(*θ*) to estimate the posterior. We chose truncated normal distributions for all priors because they allow for weighting of more plausible parameters (in contrast to e.g. uniform distributions), while they enable restrictions to plausible ranges (in contrast to e.g. normal distributions). A *n*-dimensional truncated normal distribution *𝒩*_*T*_ is defined by a mean *µ* = (*µ*_0_, *…,µ*_*n*_)^*T*^, a covariance matrix *σ* and *n*-dimensional space *W* = [*a*_1_, *b*_1_] × … × [*a*_*n*_, *b*_*n*_]:

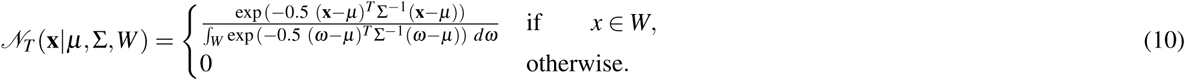

The prior means *µ*_*i*_ and truncation bounds [*a*_*i*_, *b*_*i*_] were based on experimental data wherever possible, including data from rat and different cell types such as rod bipolar cells, as well as pilot simulations. For parameter inference, we normalized the parameter space such that the truncation bounds were [0, 1] in all dimensions. The standard deviation of all priors was set to 0.3, and covariances were set to zero. To sample from *𝒩* _*T*_, we implemented a rejection sampler, that samples from a normal distribution with the same mean *µ* and covariance matrix *Σ* and resamples all **x** not in *W*.

#### Inference algorithm

SNPE estimates a posterior parameter distribution represented by a mixture-density network. Inference is performed in several rounds. In every round *i*, the algorithm samples from a proposal prior 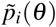 and estimates a posterior distribution *p*(***θ*|x**^0^) where **x**^0^ is a summary statistic of target data. In the first round, the proposal prior over all *P* parameters must be defined. This first proposal prior 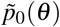 is equal to the prior *p*(*θ*). In subsequent rounds, the proposal prior is updated and differs from the prior. The prior is however still relevant for updating the posterior (see below). In every round, the algorithm draws *N* parameter samples 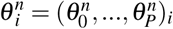 from the proposal prior 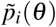. Then the multicompartment model is evaluated for all *N* parameter samples. From each simulated response, a summary statistic 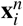 is computed, resulting in *N* pairs of parameters and summary statistics 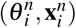. At the end of the round, a mixture-density network is trained with the summary statistics as input, and the parameters *φ* of a parametrized auxiliary Mixture of Gaussian distribution *q*_*ϕ*_ as output. The network is trained by minimizing the loss function *ℒ* :

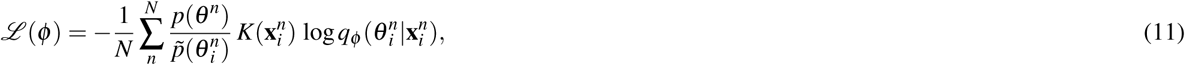

where *K* is a kernel function between zero and one that weights the influence of samples on the network training. *K* is close to one for samples with summary statistics 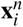 close to the the target summary statistic **x**^0^ and becomes smaller with increasing discrepancy between 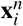 and **x**^0^. After training the network, it is evaluated at a given summary statistic **x**\* to obtain the posterior parameter distributions for the given summary statistic. Choosing **x**\* = **x**^0^ yields an approximate of the true posterior distribution *p*(*θ*|**x**^0^) ≈ *q*_*ϕ*_ (**x**^0^). This posterior can either be used as the prior for the next round, or - if the algorithm is stopped - as the final posterior distribution. A detailed proof that this actually yields an approximation of the true posterior in the Bayesian sense can be found in ^16^. We based our algorithm on the Python code available at https://github.com/mackelab/delfi version 0.5.1 with the following settings and modifications: We used truncated normal distributions with the same truncation bounds for priors and posteriors. We used only a single component for the posteriors, since we noticed that multiple components virtually always collapsed to a single component after only a few rounds of training. Instead of a multi-dimensional summary statistic we used a scalar measure of discrepancy between samples and a target (**x** can therefore be written as *x*). In the unmodified algorithm, *x*\* would therefore be zero because *x*^0^ is zero. Considering the noise in the target data, observing a discrepancy of zero is virtually impossible. Therefore, evaluating the network at *x*\* is based on extrapolation (in contrast to interpolation) which, as we observed during pilot experiments, often led to posterior estimates of poor quality or endless loops of resampling. So instead of evaluating the network at *x*^0^ = 0 in every round *i*, the network was evaluated at a the sample with the smallest discrepancy observed during this round 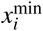. This is roughly equivalent to assuming that the best strategy for extrapolation is to simply use the estimate at the boundary. For *K*, we used a Gaussian kernel with a mean 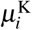 and bandwidth 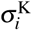 that was updated in every round *i* before network training:

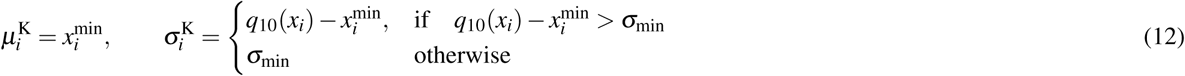

where *q*_10_(*x*_*i*_) is the 10^*th*^ percentile of all sampled discrepancies of the same round and *σ*_min_ is a lower bound for the bandwidth. *σ*_min_ was set to 0.5 if not stated otherwise.

For every neuron model, we drew 2, 000 samples per round and stopped the algorithm after the fourth round. 200 additional samples were drawn from the posterior for further analysis. For the BC models, the number of compartments was increased in this last step to 97 and 101 for the OFF- and ON-BC, respectively, equal to the number of compartments used to simulate electrical stimulation.

#### Target data of neuron models

As targets, we used two-photon imaging data recorded with an intensity-based glutamate-sensing fluorescent reporter (iGluSnFR) ^62^. All animal procedures were approved by the governmental review board (Regierungspräsidium Tübingen, Baden-Württemberg, Konrad-Adenauer-Str. 20, 72072 Tübingen, Germany) and performed according to the laws governing animal experimentation issued by the German Government. For the cone models, we used mean glutamate traces of two cone axon terminals in response to a full-field chirp light stimulus (Fig. 5). Traces were recorded in one transgenic mouse (B6;129S6-Chat^tm2(cre)Lowl^J, JAX 006410, crossbred with Gt(ROSA)26Sor^tm9(CAG-tdTomato)Hze^, JAX 007905) that expressed the glutamate biosensor iGluSnFR ubiquitously across all retinal layers after intravitreal injection of the viral vector AAV2.7m8.hSyb.iGluSnFR (provided by D. Dalkara, Institut de la Vision, Paris). Cone glutamate release in the outer plexiform layer was recorded in x-z scans (64×56 pixels at 11.16 Hz;^63^). Regions-of-interest (ROIs) were drawn manually and traces of single ROIs were then normalized and upsampled to 500 Hz as described previously^19, 64^.

For the BC models, we used mean glutamate traces of BC3a (n=19 ROIs) and BC5o (n=13 ROIs) in response to a chirp light stimulus from a recently published dataset^19^ (Fig. 6). In that study, glutamate responses were recorded from BC terminals at different depths of the inner plexiform layer (x-y scans, 64×16 pixels at 31.25 Hz). ROIs were drawn automatically based on local image correlation and traces of single ROIs were normalized and upsampled to 500 Hz (see above). Since we simulated isolated BCs (except for the cone input), we used the responses to a local “chirp” light stimulus recorded with the glycine receptor blocker strychnine, which means that the target data is less affected by inhibition from small-field amacrine cells (ACs). We did not consider input from GABAergic, wide-field ACs, because these are not strongly activated by the local chirp stimulus^19^. The shape of the BC stimulus differed from the cone stimulus as contrast was not linearized for the BC recordings and therefore intensity modulations below 20% brightness were weakly rectified.

#### Light stimulus and cell response

For a meaningful comparison between simulations and experimental data, we first matched the experimental with the simulated stimulus. For this, we linearly transformed the light stimulus such that the simulated photon absorption rates were 1×10^3^ P\**/*s for the lowest and 22×10^3^ P\**/*s for the highest stimulus intensity plus an additional background illumination causing 10×10^3^ P\**/*s, approximating the values reported in ^19^.

In NeuronC, the photon absorption rate acts as input to a phototransduction model^65^, which provides the hyperpolarizing current entering the inner segment. The membrane potential in the axon terminal compartment regulates the calcium influx into the cell which in turn influences the glutamate release rate. This glutamate release from the simulated cones modifies the opening probability (the fraction of open channels in the deterministic case) of postsynaptic receptors, which drive the BC models.

#### Discrepancy function

For every model evaluation, we computed a discrepancy value between simulated and experimental data. Since the target traces are recorded as relative fluorescence intensities, the absolute number of released glutamate vesicles can not be directly inferred from the target data, such that it only constrains relative variations in the release rate during simulation. Since we also wanted to constrain our models to plausible membrane potentials and release rates, we combined the following seven discrepancy measures:

- *δ*_iGluSnFR_: The discrepancy between the experimental and simulated iGluSnFR trace.
- 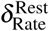: A penalty for implausibly high resting release rates.
- 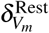: A penalty for implausibly low or implausible high resting membrane potentials.
- 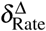: A penalty for implausibly low release rate changes.
- 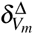: A penalty for implausibly low membrane potential changes.
- 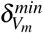: A penalty for implausibly low membrane potentials.
- 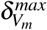: A penalty for implausibly high membrane potentials.

The total discrepancy was computed as the sum of all discrepancy measures, with all measures defined to be between zero and one.

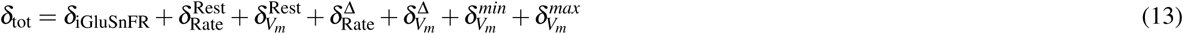

*δ*_iGluSnFR_ computes the discrepancy between an iGluSnFR target **x**_*t*_ and a simulated iGluSnFR trace **x**_*s*_. To estimate the simulated iGluSnFR trace, we convolved the glutamate release rate **r**_*s*_ with an iGluSnFR kernel *κ*. Here, *κ* was approximated with an exponential function, based on iGluSnFR intensity changes to spontaneous vesicle release reported in ^62^.

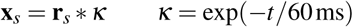

The discrepancy was then computed as the euclidean distance between the simulated and the target iGluSnFR trace with respect to a distance minimizing linear transformation of the simulated trace. This linear transformation was necessary because the target traces only reflect relative fluorescence changes. The discrepancy was normalized to be between zero and one by dividing by the variance ‖**x**_*t*_ − *µ*(**x**_*t*_)‖^2^, where *µ* is the mean, of the target data.

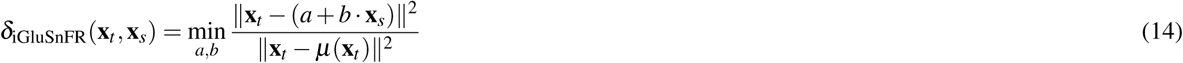

For all other discrepancies, specific values of the glutamate release rate (in the case of the BCs, the mean release rate over all synapses) or the somatic membrane potential were compared to a lower and an upper bound of target values *t*_*l*_ and *t*_*u*_, such that values within these bounds were assigned a penalty of 0.0. To constrain the discrepancy of single discrepancy measures, values outside the bounds *p*_*l*_ and *p*_*u*_ were assigned the highest penalty of 1.0. Given a specific value of *x*_*s*_ the respective discrepancy *δ*(*x*_*s*_) is computed as:

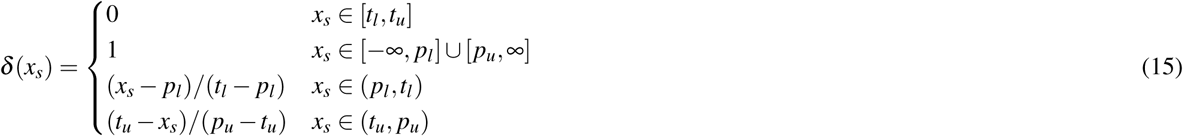

To compute 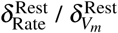 the resting release rate 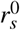 / the resting membrane potential 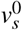 for the background light adapted state were extracted. For the BC models, the resting membrane potential was not penalized for values between *t*_*l*_ = −65 mV and *t*_*u*_ = −45 mV based on reported values for mice^30, 46^ and rat retina^42^. For the cone model, the expected resting membrane potential was more depolarized between *t*_*l*_ = −55 mV and *t*_*u*_ = −40 mV^66^.

The discrepancy of the resting release rate 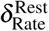 was computed similarly. For the BC models, the lower bound *t*_*l*_ was set to zero. As mentioned earlier, we limited our BC models to have a maximum sustainable release rate of 8 vesicles per second based on ^60^. We allowed non-zero resting release rates due to the background light and spontaneous vesicle fusion but constrained it to values lower than the maximum sustainable release rate^70, 71^. For the OFF-BC we chose an upper bound of 4 vesicles per second (half the maximum sustainable release rate from ^60^). For the ON-BC we chose a slightly smaller value of 3 vesicles per second. This difference was based on the observation that the ON-BC target never falls significantly below the value of the resting state, indicating that the resting release rate is probably close to zero and can therefore not become smaller. In contrast, the OFF-BC target falls below the resting value right after stimulus onset, indicating a small but non-zero resting release rate. For the cone model, we assumed a comparably high resting release rate between *t*_*l*_ = 50 and *t*_*u*_ = 80 vesicles per second based on the assumed higher maximum sustainable release rate and the fact that cones show steady release in darkness^67, 68^.

For the penalty on implausible release changes 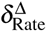 we computed the largest absolute difference Δ*r* between the resting release rate 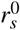 and release rates **r**_*s*_ after stimulus onset. 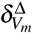 was computed analogously but for the membrane potential **v**_*s*_ and the resting membrane potential 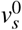:

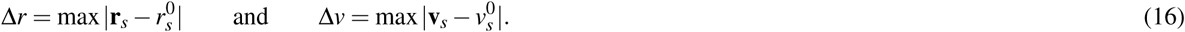

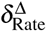 and 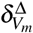 were then computed by using the differences *x*_*s*_ = Δ*r* and *x*_*s*_ = Δ*v*, respectively, in equation 15. For the BC release rate, we did not penalize differences larger than *t*_*l*_ = 5 vesicles per second. For the cone, we expected much larger differences between *t*_*l*_ = 50 to *t*_*u*_ = 65 vesicles per second due to their larger maximum sustainable release rate. For the membrane potential, we expected a difference of at least *t*_*l*_ = 5 mV based on light step responses recorded with patch clamp in mouse BCs^30, 46^. Since here, the stimulus contrast was higher, we only used the reported values as lower bounds but allowed the model to have larger variation, namely up to *t*_*u*_ = 25 mV for the OFF- and *t*_*u*_ = 15 mV for the ON-BC. We allowed greater membrane potential variation in the OFF-BC, because it receives input from more cones.

For the discrepancy measures 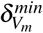 and 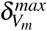 we computed the minimum and maximum of the membrane potential **v**_**s**_ after stimulus onset and used again equation 15. For 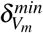 we chose *t*_*l*_ = −80 mV for the BCs and *t*_*l*_ = −60 mV for the cone model and in both cases *t*_*u*_ = ∞. For 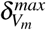 we chose *t*_*u*_ = −10 mV for the BCs and *t*_*u*_ = −35 mV for the cone, in both cases we set *t*_*l*_ = −∞. The BC values are based on data from rat^42^ and ground squirrel^69^; the cone values are based on ^66^.

All values for *p*_*l*_ and *p*_*u*_ were based on pilot simulations with the goal to distribute the penalties where they most mattered. All discrepancies (except for *δ*_iGluSnFR_) and their respective values *p*_*l*_, *p*_*u*_, *t*_*l*_ and *t*_*u*_ are illustrated in Fig. 2 for clarity.

Some parameter combinations caused the simulation to become numerically unstable. If a simulation could not successfully terminate for this reason, we set every discrepancy measure to 1.0 (the maximum) resulting in a total discrepancy of 7.0. In other cases, the BC models had a second, strongly depolarized and therefore biologically implausible equilibrium state. To test for this, we simulated a somatic voltage clamp to 30 mV for 100 ms and checked whether the membrane potential would recover to a value of −30 mV or lower within additional 300 ms. Samples not recovering to −30 mV or lower were assigned a total discrepancy of 7.0. All samples with a discrepancy of 7.0 were ignored during training of the mixture-density network.

### 2.3 Data analysis of simulated traces

For data analysis in Fig. 4, equation 14 was not only used to compute the discrepancy between simulations and the respective targets, but also more generally to compare different experimental and simulated iGluSnFR traces. For this, **x**_*s*_ does therefore not necessarily refer to a simulated trace and **x**_*t*_ not necessarily to the target used for parameter inference.

To quantify the timing precision of our neuron models, we estimated peak times in simulated and target iGluSnFR traces to compute pairwise peak time differences. For every peak in the simulated trace we computed the time difference to the closest peak of the same polarity (positive or negative) in the target resulting in approximately 50 positive and negative peaks time differences per trace.

### 2.4 Simulation of electrical stimulation

To simulate external electrical stimulation of our BC models, we implemented a two-step procedure. In the first step, the electrical field is estimated as a function of space and time across the whole retina for a given stimulation current. By setting a position of the BC multicompartment models within the retina, the extracellular voltage for every compartment can be extracted. In the second step, the extracellular voltages are applied to the respective compartments (Fig. 1C) to simulate the neural response in NeuronC. To be able to perform the first step, we estimated the electrical properties of retinal tissue first. For this we utilized the same algorithm that was used for parameter inference of the neuron models. To validate the framework, we simulated the electrical stimulation in ^72^ and compared experimental and simulated neural responses. Finally, we utilized the framework to find electrical stimuli for selective stimulation of OFF- and ON-bipolar cells. Details of the implementation and the experimental data are described in the following.

#### Computing the extracellular voltage

We estimated the electrical field in the retina for a given electrical stimulus with the finite-element method using the software COMSOL Multiphysics ®^73^. We modeled the photoreceptor degenerated retina as a cylinder with a radius of 2mm and a height of 105 µm^74^. The stimulation electrodes were modeled as flat disks on the bottom of the retina. Above the retina, an additional cylinder with the same radius and a height of 2mm was placed to model the electrolyte. The top of this cylinder was assumed to be the return electrode. The implementation of such a model with the subdivision into finite elements is shown in Fig. 3. For a single circular stimulation electrode, the model was radially symmetric and could therefore be reduced to a half cross-section as shown in Fig. 3 to reduce the simulation speed without altering the results. The following initial and boundary conditions were applied to the model. The initial voltage was set to zero at every point *V*(*x, y, z, t* = 0) = 0. The surface normal current density 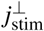 of stimulation electrodes was always spatially homogeneous and dependent on the total stimulation current *i*_stim_ and the total surface area of all electrodes *A*_electrode_:

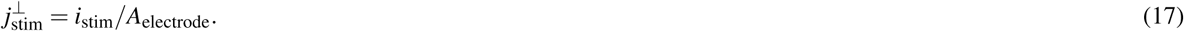

The potential of the return electrode was kept constant *V*(*t*)_return_ = 0. At all other boundaries, the model was assumed to be a perfect insulator 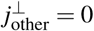. We assumed a spatially and temporally homogeneous conductivity and permittivity in both the retina and the electrolyte. The conductivity of the electrolyte was set to *σ*_ames_ = 1.54 S*/*m based on ^75^ and its relative permittivity was assumed to be *ε*_ames_ = 78, based on the value for water. The conductivity *σ*_retina_ and relative permittivity *ε*_retina_ of the retina were optimized with respect to experimental target data as described below.

#### Target data to infer the electrical parameters of the retina

To estimate the electrical properties of the retina, we first recorded target data. All procedures were approved by the governmental review board (Regierungspräsidium Tübingen, Baden-Württemberg, Konrad-Adenauer-Str. 20, 72072 Tübingen, Germany, AZ 35/9185.82-7) and performed according to the laws governing animal experimentation issued by the German Government. We applied different sinusoidal stimulation voltages *v*_stim_ and recorded the evoked currents. Currents were recorded with 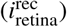 and without 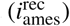 retinal tissue placed on the micro-electrode array. In both cases the recording chamber was filled with an electrolyte (Ames’ medium, A 1420, Sigma, Germany). A single Ag/AgCl pellet (E201ML, Science Products) was used as counter electrode and located approximately 1 cm above a customized micro-electrode array. The electrodes, made of sputtered iridium oxide had diameters of 30 µm and center-to-center distance of 70 µm. The stimulation current was calculated from the voltage drop across a serial 10Ω resistor in series with the Ag/AgCl electrode^72^. The voltage drop was amplified using a commercial voltage amplifier (DLPVA, Femto Messtechnik GmbH, Berlin, Germany) and recorded using the analog input (ME 2100, Multi Channel Systems MCS GmnH, Germany). Stimulation currents were measured across an ex vivo retina of a rd10 mouse (female; post-natal day 114; strain: Pde6brd10, JAX Stock No: 004297).

We applied sinusoidal voltages of 25 and 40 Hz. For 25 Hz, we applied amplitudes from 100 to 600mV with steps of 100mV. For 40Hz all amplitudes were halved. For both frequencies, two of the seven applied amplitudes are shown in Fig. 8B.

**Figure 7.**
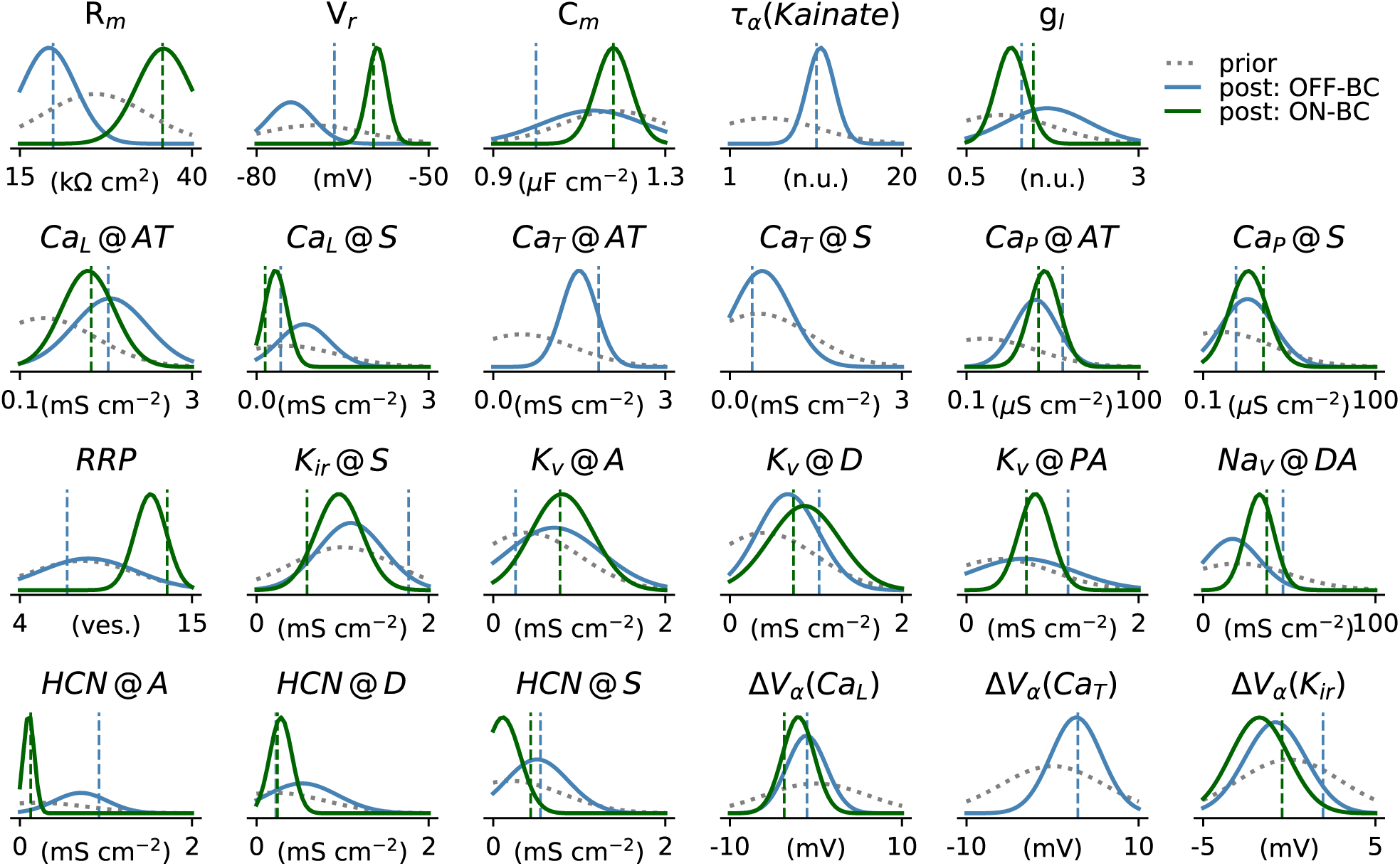
Parameter distributions of the BC models. 1D-marginal prior (dashed gray line) and posterior distributions (solid lines) are shown for the OFF- (blue) and ON-BC (green). The parameters of the posterior samples with the lowest total discrepancy are shown as dashed vertical lines in the respective color. X_Y_ @Z refers to the channel density of channel X_Y_ at location Z. Locations are abbreviated; S: soma, A: axon, D: dendrite and AT: axon terminals (see Fig. 1 and main text for details). Note that although these 1D-marginal distributions seem relatively wide in some cases, the full high-dimensional posterior has much more structure than a posterior distribution obtained from assuming independent marginals (see Fig. 4). Not all parameter distributions are shown.

**Figure 8.**
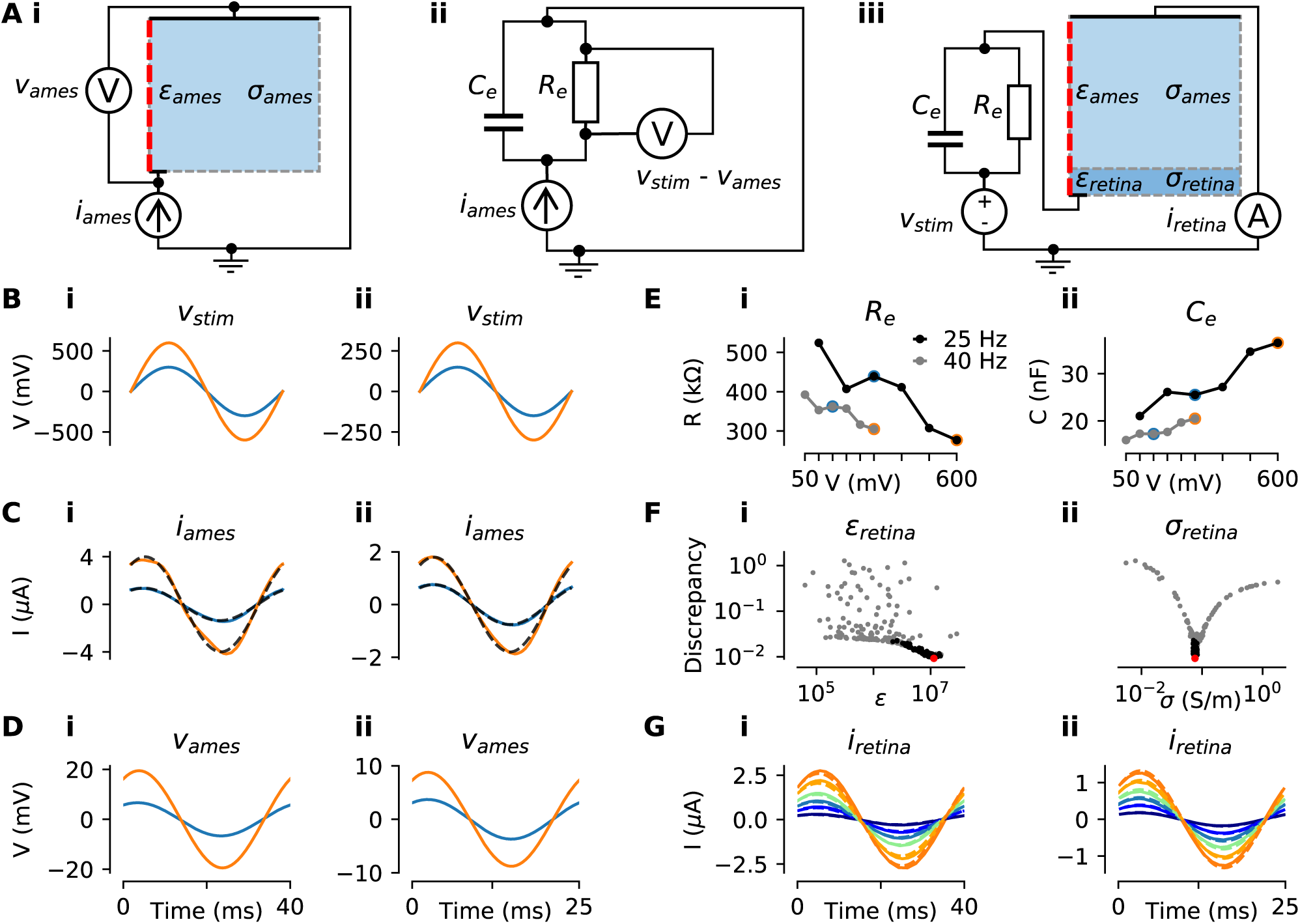
*Estimation of the conductivity σ*_*retina*_ *and the relative permittivity ε*_*retina*_ *of the retina for simulating electrical stimulation*. ***(A)*** *Electrical circuits used during parameter estimation*. ***(B)*** *Stimulation voltages v*_*stim*_ *at* 25 (*left) and* 40 Hz *(right). From the six experimentally applied amplitudes, only the amplitudes used for parameter inference are shown*. ***(C)*** *Experimentally measured currents i*_*ames*_ *through electrolyte (Ames’ solution) without retinal tissue for the stimulus voltages v*_*stim*_ *in (B). The mean traces over all (but the first two) repetitions are shown (black dashed lines). Sine waves were fitted to the mean traces (solid lines) with colors referring to the voltages in (B)*. ***(D)*** *Simulated voltages over the electrolyte v*_*ames*_ *using the fitted currents in (C) as stimuli applied to the circuit in (Ai)*. ***(Aii)*** *Electrical circuit used to model the electrode plus interface*. ***(E)*** *Stimulus frequency and amplitude dependent estimates of R*_*e*_ *(i) and C*_*e*_ *(ii) based on the electrical circuit shown in (Aii) for* 25 (*black) and* 40 Hz *(gray). Note, that the values were derived analytically (see main text). The values corresponding to the stimulus voltages shown in (B) are highlighted with respective color*. ***(Aiii)*** *Electrical circuit used to estimate σ*_*retina*_ *and ε*_*retina*_. *The respective values for R*_*e*_ *and C*_*e*_ *are shown in (E) and are dependent on v*_*stim*_. *The current i*_*retina*_ *through the model is measured for a given stimulus voltage v*_*stim*_. ***(F)*** *Sampled parameters of ε*_*retina*_ *and σ*_*retina*_ *and the respective sample losses. First, samples were drawn in a wide logarithmic space (gray dots) and then in a narrower linear parameter space. The best sample (lowest discrepancy) is highlighted in red*. ***(G)*** *Simulated currents i*_*retina*_ *(solid lines) through the circuit in (Aiii) with optimized parameters (red dot in (E)) and respective experimentally measured currents (broken lines). Here, results for all six stimulus amplitudes are shown for both frequencies*.

#### Procedure to infer the electrical parameters of the retina

We estimated the conductivity *σ*_retina_ and relative permittivity *ε*_retina_ of the retina in three steps based on the experimental voltages *v*_stim_ and the respective recorded currents 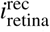 and 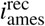. To facilitate the following steps we fitted sinusoids *i*_retina_ and *i*_ames_ to the slightly skewed recorded currents and used them in the following (Fig. 8C). To fit the sinusoids, we minimized the mean squared error between recorded currents and idealized sinusoidal currents of the same frequency *f*, resulting in estimates of the phase *ϕ*(*i*_ames_) and the amplitude *A*(*i*_ames_) of the currents:

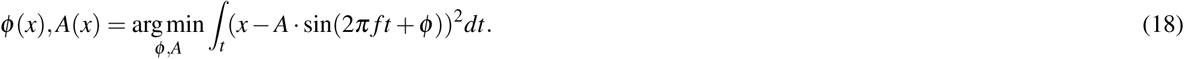

During parameter inference, we only used two voltage amplitudes per frequency, resulting in four voltage and eight current traces. The other amplitudes were used for model validation. First, we estimated the electrical properties of the electrode. Here, “electrode” is meant to include the electrical double layer and all parasitic resistances and capacitances in the electrical circuit. We simulated the voltage *v*_ames_ across the electrolyte without retinal tissue by applying the currents *i*_ames_ as stimuli (Fig. 8Ai). Since this setup does not contain anything besides the electrolyte and the electrode, the difference between the experimental stimulus *v*_stim_, which was applied to record *i*_ames_, and the simulated voltage *v*_ames_ was assumed to have dropped over the electrode:

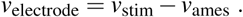

Based on that assumption we could estimate the electrical properties of the electrode. We modeled the electrode as a *RC* parallel circuit (Fig. 8Aii). Having both, sinusoidal voltages (*v*_electrode_) over and the respective sinusoidal currents (*i*_ames_) through the electrode, we analytically computed the values for *R*_*e*_ and *C*_*e*_ as follows. We assumed *R*_*e*_ and *C*_*e*_ to be dependent on *v*_stim_ and therefore to be dependent on the stimulus frequency and amplitude. From the data we derived the phase *ϕ*_*Z*_ and amplitude |*Z*| of the impedance formed by the *RC* circuit. For every *v*_electrode_, we estimated *ϕ*(*v*_electrode_) and *A*(*v*_electrode_) using equation 18. *ϕ*_*Z*_ and |*Z*| were then computed as:

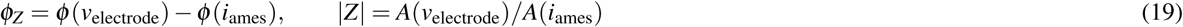

Then, knowing the frequency *f, ϕ*_*Z*_ and |*Z*| are sufficient to compute *R* and *C*:

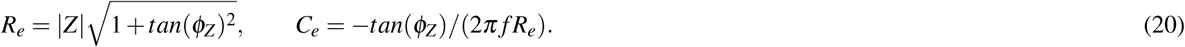

With the estimated values of the *RC* circuit, we created a model with only two unknowns, the conductivity *σ*_retina_ and the relative permittivity *ε*_retina_ of the retina (Fig. 8Aiii). To estimate the unknown parameters of this model, we used the same inference algorithm as for the neuron models but with a different discrepancy function. Here, the discrepancy *δ*_*R*_(*v*_stim_) for a stimulus *v*_stim_ was computed as the mean squared error between the respective experimental current (now with retinal tissue) *i*_retina_ and the simulated current 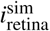:

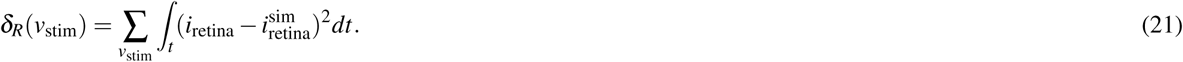

The total discrepancy was computed as the sum of all discrepancies *δ*_*R*_(*v*_stim_) for the four different *v*_stim_ stimuli that were used. To cover a wider range of possible parameters, we first estimated the parameters in a logarithmic space by sampling the exponents *p*_*σ*_ and *p*_*ε*_ of the parameters:

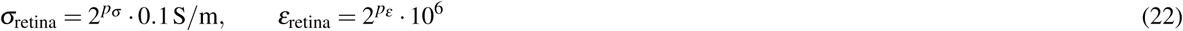

We used normal distributions (without truncation) as priors for *p*_*σ*_ and *p*_*ε*_ and set the means to 1.0 and the standard deviations to 2.0. After three rounds with 50 samples each, we computed the minimum (*a*_*σ*_, *a*_*ε*_), maximum (*b*_*σ*_, *b*_*ε*_) and mean (*µ*_*σ*_, *µ*_*ε*_) for both parameters *σ*_retina_ and *ε*_retina_ from the 10% best samples. Then, we then ran the parameter inference algorithm again, but now in a linear parameter space around the best samples observed in the logarithmic space. For the priors of *σ*_retina_ and *ε*_retina_ we used truncated normal priors bound to [*a*_*σ*_, *b*_*σ*_] and [*a*_*ε*_, *b*_*ε*_] respectively with means *µ*_*σ*_ and *µ*_*ε*_ and standard deviations of 0.3. The parameters resulting in the lowest sampled discrepancy during optimization are referred to as the optimized parameters and were used to simulate the neural responses to electrical stimulation.

#### Simulation of the neural response to electrical stimulation

With the optimized parameters for the electrical properties of the retina, we were able to compute the BC responses for any given stimulation current. Note that for this, we used the model illustrated in Fig. 3 as described earlier but with the optimized parameters for *σ*_retina_ and *ε*_retina_. To simulate the neural response, we first used the stimulation current to simulate the extracellular voltage over time within the retina. After defining the relative position of the multicompartment model with respect to the retinal cylinder, we extracted the extracellular voltage for each compartment at its the central position (Fig. 3C). Finally these extracellular voltages were applied to the compartment models in NeuronC to simulate their response (Fig. 1C). In all simulations, we modeled subretinal stimulation of photoreceptor degenerated retina^76^. For this, we removed all cone input from the BCs and virtually placed the multicompartment models in the retinal cylinder such that the dendrites were facing towards the electrode. The z-position of BC somata, i.e. the distance to the bottom of the retinal cylinder, was set to 30 µm.

#### Model validation

To validate the model for electrical stimulation, we compared simulated BC responses to experimentally recorded retinal ganglion cell (RGC) thresholds to 4ms biphasic current pulses reported in ^72^. In this study, the RGC thresholds were recorded epiretinally under subretinal stimulation of photoreceptor degenerated (rd10) mouse retina using a micro-electrode array (Fig. 3A). The stimulation threshold was defined as the charge delivered during the anodic stimulation phase evoking 50% of the firing rate of a specific RGC. On the micro-electrode array. The 30 µm diameter electrodes were arranged on a regular grid with a center-center spacing of 70 µm. The RGC thresholds were measured for different numbers *N* of *N*×*N* active electrodes.

We simulated the electrical field in the retina for the configurations with 1×1, 2×2, 4×4 and 10×10 active electrodes using the respective currents from the experimental data. The electrodes were centered with respect to the retinal cylinder. For every stimulation current, we simulated the response of the OFF- and ON-BC at 20 equidistant xy-positions with a distance between 0 and 671 µm relative to the center (Fig. 9A). Simulation temperature *T*_sim_ was set to 33.5 °C to match experimental conditions. Simulations were 30 ms long, beginning 1ms before stimulus onset. For every BC and stimulus, we computed the mean number of vesicles released per synapse.

**Figure 9.**
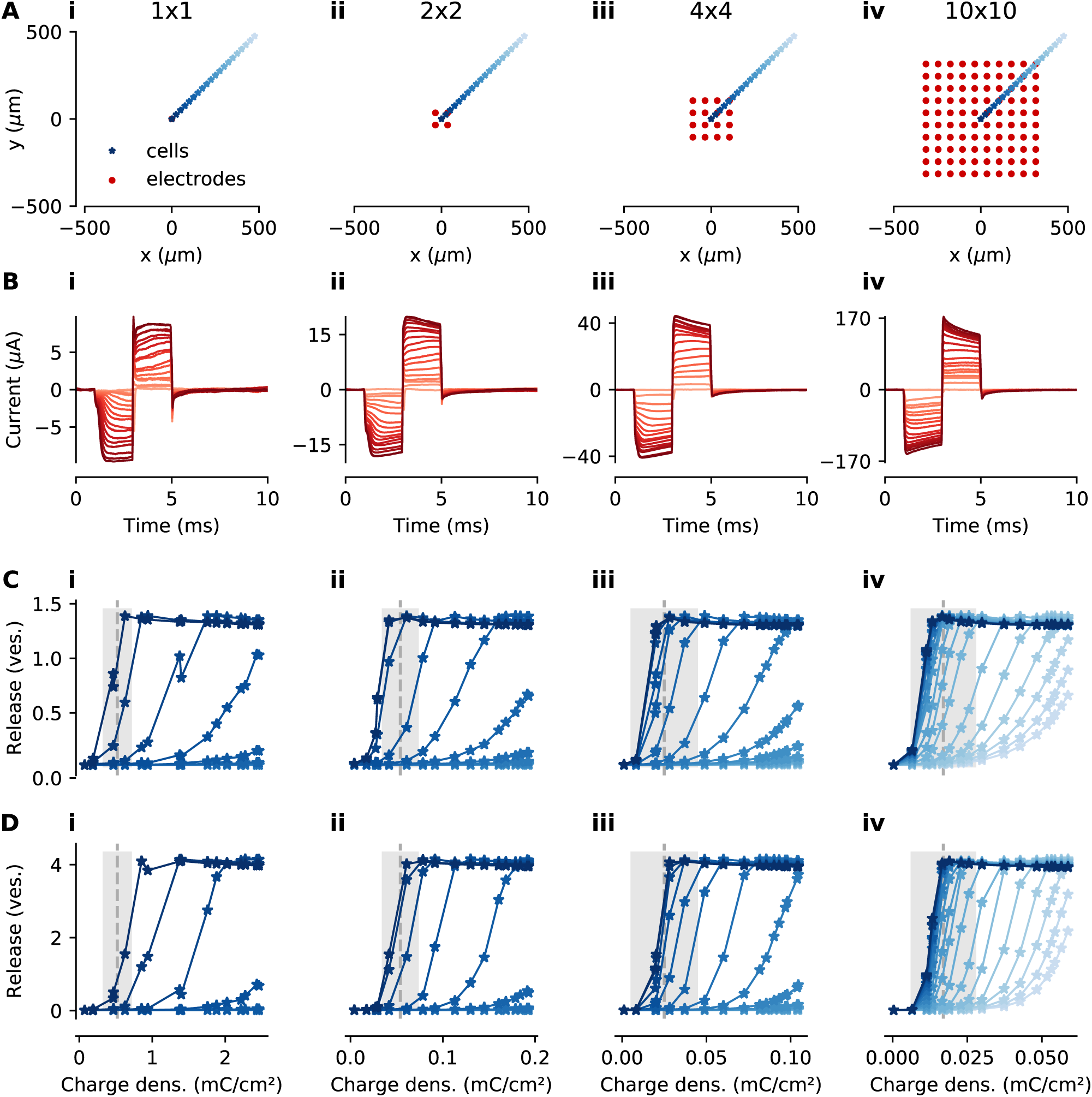
*Threshold of electrical stimulation for experimentally measured RGCs and simulated BCs of photoreceptor degenerated mouse retina*. ***(A)*** *xy-positions of BCs (blue stars) and electrodes (red dots) for* 1×1, 2×2, 4×4 *and* 10×10 *stimulation electrodes, respectively. Every electrode is modeled as a disc with a* 15 µm *radius. Except for the electrode configuration, the models were as in Fig. 8*. ***(B)*** *Stimulation currents measured experimentally and used as stimuli in the simulations*. ***(C, D)*** *Glutamate release (mean over all synapses) of simulated OFF- and ON-BCs respectively. Glutamate release is shown for different charge densities (x-axis) and cell positions (shades of blue correspond xy-positions in (A); e*.*g. the darkest blue corresponds to the central BC). Experimentally measured RGC thresholds (gray dashed lines) plus and minus one standard deviation (gray shaded ares) are shown in the same plots*.

#### Optimizing electrical stimulation to separately activate ON- and OFF-BCs

To find stimuli for selective stimulation of ON- and OFF-BCs, we simulated the response of the BC models to different electrical stimuli. For this, we used a single 30 µm diameter electrode and centered the dendrites of the simulated BCs above this electrode (Fig. 10E,F). To find stimuli that stimulate the OFF-BC without stimulating the ON-BC or vice versa, we utilized the same algorithm used for estimating the BC parameters. Here, the inference algorithm was used to estimate parameters of a 40 ms stimulation current *i*_stim_ parametrized by four free parameters *p*_1_, …, *p*_4_. The current was defined as a cubic spline fit through the knot vector **a** = (0, *p*_1_, …, *p*_4_, *p*\*, 0) spaced equidistantly in time between zero and 40 ms, where *p*\* is chosen such that the stimulus is charge neutral (i.e. the integral over the current is zero). For all stimuli, the maximum stimulus amplitude was normalized to 0.5 µA. An illustration is shown in Fig. 10.

**Figure 10.**
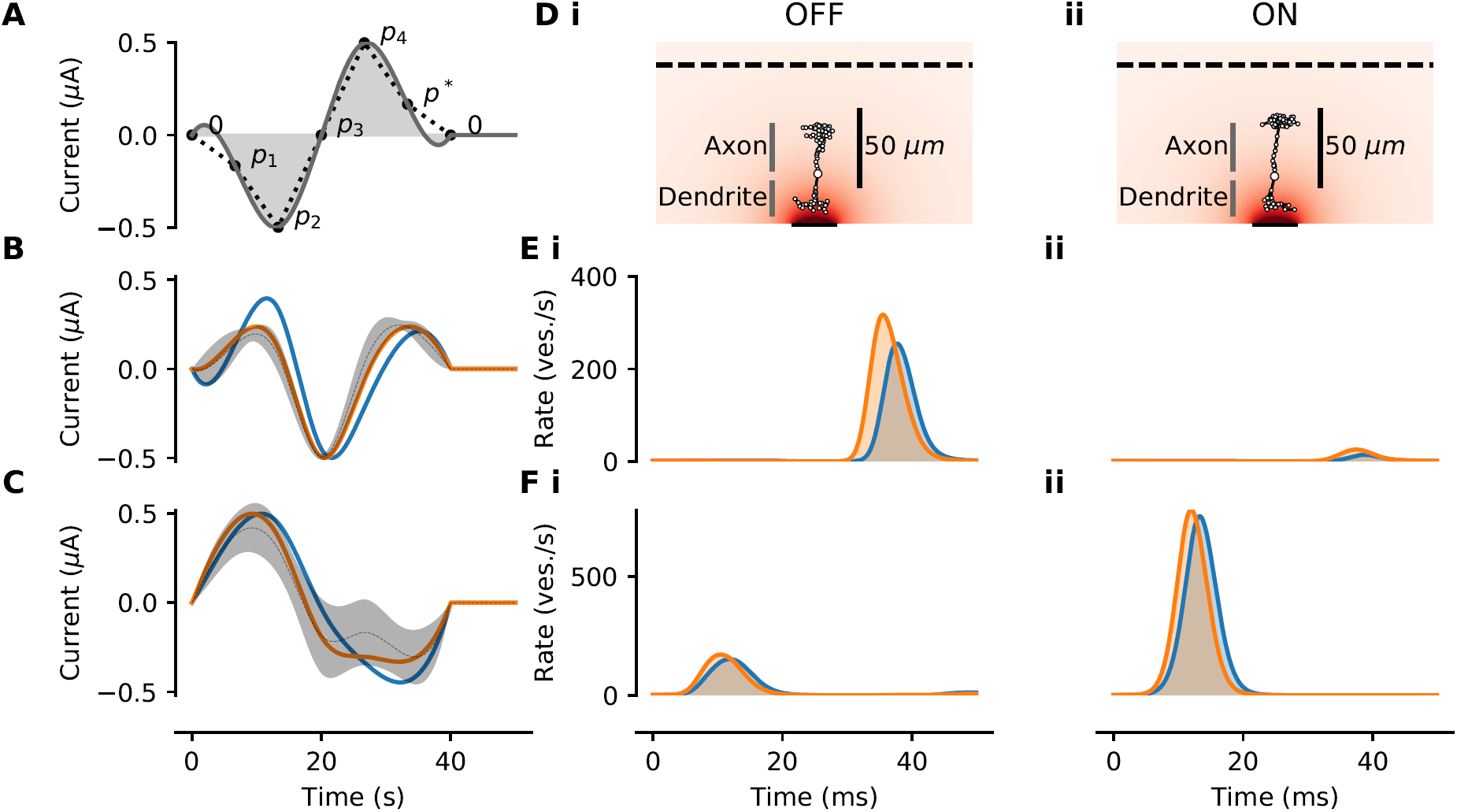
*Electrical currents optimized for selective ON- and OFF-BC stimulation*. ***(A)*** *Illustration of the stimulus parametrization. The stimulus is dependent on four free parameters p*_1_…*p*_4_ *that define the relative current amplitudes over time. p*\* *is always chosen such that the stimulus is charge neutral, i*.*e. the integral over the current (gray shaded area) is zero. The stimulation current is computed by fitting a cubic spline through points defined by p*_1_…*p*_4_ *and p*\* *that are placed equidistantly in time (black dots). The currents are normalized such that the maximum amplitude is always* 0.5 µA. ***(B, C)*** *Optimized stimulus distribution and example stimuli for selective OFF- and ON-BC stimulation respectively. The best stimulus observed during optimization (blue) and a second example stimulus with similar quality but slightly different shape (orange) are shown. Additionally, the mean (gray dashed line) plus-minus one standard deviation (gray shaded area) from* 200 *stimuli sampled from the posterior distribution are shown*. ***(D)*** *Illustration of the electrical stimulation of an OFF- (i) and ON-BC (ii) multicompartment model*. ***(E, F)*** *Release rates as means over all synapses for the OFF- (i) and ON-BC (ii) in response to the electrical stimuli shown in (B) and (C) respectively. Stimuli and responses are shown in matching colors*.

Here, the priors over *p*_1_, …, *p*_4_ were uniform between −1 and 1. For every sampled stimulus 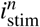, we simulated the response of the BCs for 50 ms starting with the stimulus onset. The discrepancy measure 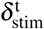 for parameter estimation was computed by comparing the mean release rate over all synapses and time of the OFF-BC 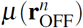 to the mean release rate of the ON-BC 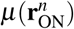. Dependent on whether the ON- or OFF-BC was defined to be the stimulation target t, the discrepancy was computed as:

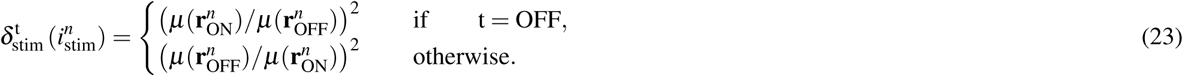

We ran the parameter inference twice, once with the ON and once with the OFF-BC as target *t*.

### Code and data availability

Models, simulation code and data will be available upon publication at https://github.com/berenslab.

## 3 Results

We used a high-resolution electronmicroscopy data set^18^ to create biophysically realistic multicompartment models of three neuron types from the mouse retina including cones, an OFF- and an ON-bipolar cell (BC) type. These neurons form the very beginning of the visual pathway, with cones converting light into electrochemical signals and providing input via sign-preserving and -reversing synapses to OFF- and ON-BCs, respectively. The parameters of these models include the basic electrical properties of the cells as well as the density of different ion channel types in different compartments. Given a set of parameters, simulations from the model can easily be generated; however, it is not possible to evaluate the likelihood for a given set of parameters, which would be required for standard parameter optimization.

To overcome the challenge of choosing the resulting 20 to 32 parameters of these models, we adapted a recently developed technique called Sequential Neural Posterior Estimation (SNPE)^16^ (for details, see Methods). Starting from prior assumptions about the parameters, the algorithm compared the result of simulations from the model to data obtained by two-photon imaging of the glutamate release from the neurons^19^ and measured a discrepancy value between the simulation and the data. Based on this information, the algorithm used a neural network to iteratively find a distribution over parameters consistent with the measured data. This yielded optimized biophysically realistic models for the considered neuron types.

### 3.1 Inference of cone parameters

We first estimated the posterior distribution over the parameters of a cone based on the glutamate release of a cone stimulated with a full-field chirp light stimulus, consisting of changes in brightness, temporal frequency and temporal contrast (Fig. 4A and 5). The cone model had a simplified morphology and consisted of four compartments (Fig. 1, see Methods). We included a number of ion channels in the model reported to exist in the cones of mice or closely related species (see Tables 1). Prior distributions were chosen based on the literature. For inference, we drew 2, 000 samples of different parameter settings from the proposal prior per round and stopped the algorithm after the fourth round. Then, 200 more parameter samples were drawn from the respective posteriors for further analysis. The chosen discrepancy functions penalized discrepancies between the target and simulated iGluSnFR trace *δ*_iGluSnFR_, implausible membrane potentials and release rates, and were added to yield a total discrepancy *δ*_tot_.

We found that the total discrepancy *δ*_tot_ of the cone model was relatively high for most samples drawn from the prior but decreased over four rounds of sampling (Fig. 4A). Almost no samples with the highest possible discrepancy were drawn after the first round. We observed that the discrepancy measuring the fit quality to the glutamate recording *δ*_iGluSnFR_ was relatively small in first few optimization rounds, before increasing again. This was caused by an increase in membrane potential variation and a large increase in the mean release rate between the third (median of mean release rates: 35) and fourth round (median of mean release rates: 72), which was necessary to reduce the rate and membrane variation discrepancies, 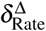 and 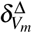.

The parameter setting with lowest discrepancy (*δ*_tot_ = 0.11) modeled accurately the response of the cone to full-field stimulation with the chirp light stimulus (Fig. 5A-D). The simulated iGluSnFR signal nicely matched the data both on a coarse timescale and in the millisecond regime (Fig. 5D). Indeed. for this sample, all discrepancies besides *δ*_iGluSnFR_ were equal to zero and most of the remaining discrepancy could be attributed to the noisy target data.

As our inference algorithm returned not only a single best set of parameters, but also a posterior distribution, we could obtain additional parameter samples from the model which should produce simulations consistent with the data. Almost all samples from the posterior yielded simulations that matched the target data well (median *δ*_iGluSnFR_: 0.12) and the overall total discrepancy was small (median *δ*_tot_: 0.43). Besides the discrepancy between the experimental and simulated glutamate trace *δ*_iGluSnFR_, most of the remaining discrepancy in the posterior samples was caused by rate variation (median 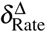: 0.11) and resting rates (median 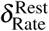: 0.00) that were too low in some of the models. While in principle we could propagate the remaining uncertainty about the model parameters provided by the posterior to the inference about BC models, we used only the parameter set with the smallest discrepancy for efficiency and refer to this as the optimized cone model.

### 3.2 Inference of bipolar cell parameters

We next turned to anatomically detailed multicompartment models of two BC types. We chose to model type 3a and type 5o because these were the OFF- and ON-CBC types for which we could gather most prior information from the literature. The anatomy of the cells was extracted from a 3D reconstruction of examples of these cell types based on electron microscopy data^18^ and divided into regions sharing ion channel parameters (Fig. 1). As for the cone model, the channels included in the model and the prior distributions were chosen based on the literature (see Table 1). This yielded 32 and 27 free parameters for the OFF- and ON-BC, respectively.

We fitted the BC type models to published two-photon glutamate imaging data reflecting the glutamate release from the BC axon terminals^19^. In this case, we used responses to a local chirp light stimulus activating largely the excitatory center of the cells. In addition, the responses were measured in the presence of the drug strychnine to block locally acting inhibitory feedback through small-field ACs^19^ (see Methods for details). Similar to what we observed in cones, the total discrepancy *δ*_tot_ for parameter sets sampled for the OFF- and ON-BC model decreased over the four rounds of optimization (Fig. 4B and C). In contrast to what we observed for the cone model, the discrepancy term penalizing deviations from the glutamate trace *δ*_iGluSnFR_ declined approximately equally fast as the total discrepancy *δ*_tot_.

We found that simulations generated with the parameter set with minimal total discrepancy or parameters sampled from the posterior matched the target traces very well for both OFF- and ON-BC models (Fig. 6). As for the optimized cone model, the optimized BC models, e.g. the BC samples with the lowest total discrepancies, had discrepancies of zero except for the iGluSnFR discrepancy *δ*_iGluSnFR_. To get a more quantitative impression of the quality of the model fits we compared the pairwise iGluSnFR discrepancies *δ*_iGluSnFR_ between the optimized BC models, the experimentally measured response traces as used during optimization, traces recorded from the same cell type without application of strychnine and responses of another OFF- and ON-BC. For both optimized cell model outputs, the discrepancy was smallest for the targets used during optimization. This shows that the optimized models were able to reproduce cell-type specific details in light response properties that go beyond the simple distinction of ON and OFF responses. While the discrepancies between traces of different ON-BC types were overall relatively small for local stimulation^19^, the discrepancies between traces from OFF cells were larger likely due to network modulation of the target cell type by ACs (indicated by the difference between the target and the no-drug condition) and larger response differences between the two compared OFF-BC types. The posterior samples of both BC models had a low discrepancy, except for a few samples (median *δ*_tot_: 0.27 and 0.23 of the OFF- and ON-BC, respectively). The only discrepancy measure with a non-zero median was *δ*_iGluSnFR_ which also accounts for 74% and 79% of the mean total discrepancy for the OFF- and ON-BC respectively.

Despite the overall high resemblance between optimized model outputs and targets, there were some visible systematic differences. For the ON-BC, the target showed a skewed sinusoidal response with faster rise than fall times during the frequency and amplitude increasing sinusoidal light stimulation between 10 s and 18 s and between 20 s and 27 s respectively. In contrast, the optimized model output showed approximately equal rise and fall times, resulting in a systematic delay of positive and negative peaks (median delay of all peaks: 18 ms) in the simulated iGluSnFR trace relative to the target (Fig. 6G). Additionally, some of the positive peaks of the optimized ON-BC model during sinusoidal light stimulation were too small (e.g. at 11.5 s). This effect might have been a side-effect of the peak timing difference between target and model: Amplitude increases were inefficient in reducing the discrepancy as long as the peaks were not precisely aligned. In contrast, the peak time precision of the OFF-BC model was much higher (median delay of all peaks: 2 milliseconds). In this case, the main difficulty for the model appeared to be its inability to reproduce the non-linearity in the cell response to the increasing amplitude sinusoidal light stimulation between 20 s and 27 s.

After having verified that the posterior over parameters provided a good fit to the experimental data, we inspected the one dimensional marginal distributions to learn more about the resulting cellular models (Fig. 7). For all parameters, the marginal posteriors had smaller variances than the priors, indicating that the parameter bounds were not chosen too narrow. For some parameters, the posterior mean differed substantially from the prior mean (e.g. the T-type calcium channel density at the axon terminal of OFF-BC) while it was largely unchanged for others (e.g. the area specific membrane resistance of ON-BC). In addition, the full posterior in the high-dimensional parameter space led to simulations which were on average better (median: 0.27 vs. 0.33 and 0.23 vs. 0.36 for the OFF- and ON-BC, respectively) and less variable in their quality (95%-CIs: 0.69 vs. 1.11 and 0.56 vs. 1.15 for the OFF- and ON-BC, respectively) than parameters drawn from a posterior obtained by assuming independent marginal distributions. In most cases, the parameters resulting in the lowest total discrepancy were close to the means of the respective posteriors. For some parameters there was a strong difference between the marginal posteriors of the OFF- and ON-BC. For example, the two parameters controlling the leak conductance, *V*_*r*_ and *R*_*m*_, were much lower for the OFF-BC consistent with the strong variation of membrane resistances reported in ^22^. The membrane conductance was lower for the ON-BC, which could increase signal transduction speed in the longer axon. Even though the posteriors were narrower than the priors, they still covered a wide range of different parameters. To some extent this may reflect the fact that we fit the model parameters solely on the cells output, and e.g. dendritic parameters may be underconstrained by such data; in addition, it may also reflect variability between cells of the same type seen in the experimental data that has also been reported in other studies^19^.

After the fourth optimization round, 200 samples were drawn from the posterior distribution with an increased number of compartments to find model parameters to simulate electrical stimulation (see Methods). For comparison we also ran simulations with the same parameters, but the original number of compartments (data not shown). Interestingly, more than 90% of the samples had a lower discrepancy if the models were simulated with more compartments for both BCs. For the best 10% of the posterior samples (sorted with respect to samples with fewer compartments), all samples with more compartments had lower discrepancies. This indicates that, given enough computational power, the same parameter inference approach but with more compartments could further improve the model outputs.

### 3.3 Simulating electrical stimulation of the retina

To provide an exemplary use-case for our data-driven biophysical models of retinal neurons, we asked whether we could use our simulation framework to optimize the stimulation strategy for retinal neuroprosthetic implants. These implants have been developed for patients who lose sight due to degeneration of their photoreceptors^76^. While existing implants have been reported to provide basic vision^76–78^, they are far from perfect. For example, most current stimulation strategies likely activate OFF- and ON-pathways at the same time^79^. To this end, we created a simulation framework for subretinal electrical stimulation of retinal tissue with micro-electrode arrays. We estimated the conductivity and relative permittivity of the retina based on experimentally measured currents evoked by sinusoidal voltages and then validated simulations of the electrical stimulation of our fitted BC models with standard pulse like stimuli against responses measured in RGCs^72^. Finally, we used the simulation framework to find stimuli that can separately stimulate OFF- and ON-BCs.

Our framework for simulating the effect of external electrical stimulation using the inferred BC models consisted of two steps: we first estimated the electrical field resulting from a stimulation protocol as a function of space and time across the whole retina (Fig. 3). The corresponding extracellular voltages were then applied to the respective compartments to simulate the neural response. To do so, we needed a model of the electrical properties of the electrode and the retinal tissue. We assumed disk electrodes and a simplified model assuming homogeneous electrical properties of the retina and the surrounding medium (see Methods). This model contained two free parameters that needed to be estimated from data: the conductivity and relative permittivity of the retinal tissue.

To estimate these parameters we recorded electrical currents resulting from sinusoidal voltage stimulation with different frequencies in a dish with and without a retina (Fig. 8B, C). We used the data without a retina to estimate the properties of the stimulation electrode (Fig. 8A, D, E and Methods). Based on the estimates of the electrode properties and the data recorded with a retina, we estimated the conductivity and relative permittivity of the retina with the same parameter inference method as for the neuron models.

We found that both parameters are very well constrained by the measured data (Fig. 8F). The parameters resulting in the lowest discrepancy were *σ*_*retina*_ = 0.0764 S*/*m and *ε*_*retina*_ = 1.14*e*7 in accordance with the conductivity of 0.077 S*/*m reported for rabbit^80^ and the relative permittivity of gray matter estimated in ^81^. With these parameters, we simulated currents for all stimulus amplitudes we recorded experimentally. The simulated and experimental currents matched for the amplitudes used during parameter inference but also for amplitudes reserved for model validation (Fig. 8G). Therefore, we used them in all following experiments.

To validate our simulation framework, we compared simulated BC responses to experimentally measured RGC thresholds^72^. We simulated BCs at different positions for four different electrode configurations (Fig. 9A) and 16 stimulation current wave forms (Fig. 9B). For small stimulus charge densities, the BCs barely respond, while for very high charge densities the cells go into saturation (Fig. 9C and D). In between, the response increases from no response to saturation dependent on the distances of the simulated cell to the stimulated electrodes. Across stimulation conditions, these threshold regions coincide with the measured RGC thresholds to the same stimulation, when assuming that the stimulated RGCs were not too far away from the stimulation electrode. Since the reported RGC thresholds likely result from indirect stimulation via BCs, the consistency between the RGC and simulated BC thresholds can be interpreted as evidence that our model was well calibrated to simulate electrical stimulation.

### 3.4 Optimized electrical stimulation for selective OFF- and ON-BC stimulation

We finally used our framework for electrical stimulation to find stimuli that excite OFF- or ON-BCs selectively. To this end, we performed Bayesian inference over an electrical charge-neutral stimulus using the SNPE algorithm, using the response ratio between the two BC types as a discrepancy function to minimize (Fig. 10A). Using this procedure, we found two distinct stimulus waveform distributions able to separately stimulate OFF- and ON-BCs (Fig. 10B and C). Triphasic, anodic first stimuli evoked substantially higher neurotransmitter release in the OFF- than the ON-BC (Fig. 10B, D, E), with a cathodic middle phase of high amplitude. These stimuli were very consistent and evoked almost no response in the ON-BC (Fig. 10Eii). The stimuli optimized to target the ON-BC were all biphasic, anodic first currents (Fig. 10C, D, F). The posterior samples and the best samples during parameter inference showed little variation in the anodic phase, while the cathodic phase varies in shape. These currents targeting the ON-BC evoked a response - albeit comparably weak - of the OFF-BC (Fig. 10Fi). This resulted in a 12 times higher discrepancy for the best current targeting the ON-BC compared to the best current targeting the OFF-BC, consistent with the lower threshold for OFF-BCs to biphasic current pulses (Fig. 9).

## 4 Discussion

Here we built on recent developments in Bayesian likelihood-free inference to perform Bayesian parameter inference for complex, mechanistic models in sensory and cellular neuroscience. In particular, we showed how to use a recently developed likelihood-free parameter inference method called Sequential Neural Posterior Estimation (SNPE)^16^ to estimate the parameters of multicompartment models of retinal neurons based on light-response measurements, build a model for electrical stimulation of the retina, and optimize electrical stimulation protocols for retinal neuroprosthetic devices designed to support vision in the blind.

Performing Bayesian inference for mechanistic models is difficult, as they typically do not come with an analytical solution of the likelihood function. The SNPE algorithm — like many approximate Bayesian computation (ABC) methods^82^ — does not require such an analytical formulation, as it builds on simulations of the model. In contrast to standard ABC methods, the SNPE algorithm makes efficient use of simulation time by using all simulated data to train a mixture density network to update the parameter distributions^16, 17^. Moreover, SNPE makes minimal assumptions about the simulator, giving full flexibility to use it with different simulation environments and software. As all Bayesian methods, SNPE allows the incorporation of literature knowledge in the form of priors which can be used to constrain parameters and to put more weight on more plausible parameters. Finally, it does not only yield a point-estimate of the best parameters — like exhaustive grid-search techniques^9–11^ or evolutionary algorithms^12–15^ — but instead returns a posterior distribution that reflects remaining parameter uncertainties and allows one to detect dependencies between parameters.

Recently, there has been a surge of interest in Bayesian simulator-based inference with many recently published algorithms^16, 17, 82–89^. While we initially evaluated different algorithms, we did not perform a systematic comparison or benchmarking effort, which is beyond the scope of this project. Much of the literature on simulator-based inference evaluates the proposed algorithms on fairly simple models. In contrast, we used SNPE here to perform parameter inference of comparatively complex multicompartment models of neurons. Importantly, we did not need targeted experiments to constrain these models, but based our framework on two-photon imaging data of glutamate release in response to simple light stimuli using a genetically encoded indicator called iGluSnFR^19, 62^. This methods allows direct experimental access to the neurotransmitter release of all excitatory neurons of the retina^90^. Using this data, we inferred the distributions of model parameters relevant for all the intermediate steps between light stimulation of cones to the glutamate release from synaptic ribbons. While we optimized many parameters in the models using SNPE, we chose to keep some parameters on sensible default values to avoid issues with computational complexity. Of course, it is possible that optimization of the full parameter space would have yielded slightly better results or that some parameters would have been set to slightly different values, as a mechanism whose parameter was allowed to vary compensated for the one whose parameter was held fixed. As an alternative to our approach, one can combine classical systems identification approaches with inference for only some of the biophysical mechanisms such as the ribbon synapse^91^. Our approach, however, allows the exploration of mechanisms within neurons which are not or only barely experimentally accessible. For example, in BCs, it is currently difficult to obtain direct voltage measurements from any part of the cell but the soma. If one is interested in how the electrical signal propagates through the axon or the axon terminal, our simulations may help to obtain mechanistic insights and develop causal hypotheses.

Because our inference framework can work with experimental measurements which can be performed in a high-throughput manner, it allows for a comparably easy scaling to infer model parameters of a larger number of multicompartment models e.g. of different neuron types. In principle it could even be possible to infer the parameters of a neuron by imaging another neuron. For example, one could attempt to infer parameters of an AC by observing the neurotransmitter release of a connected BC — although such an indirect inference would likely result in larger uncertainties. Ideally, such a large scale approach would also include realistic morphologies e.g. from electron microscopy as shown here. In fact, BCs are anatomically relatively “simple” neurons, and it will be interesting to test our inference methods on neurons with more complicated structure such as some ACs^92^. While we did not aim at an exhaustive analysis of the effect of morphology on the neuron responses, one could easily explore how details of the morphology influence the distribution of optimal biophysical parameters.

Further, we extended our model to simulate and optimize external electrical stimulation of the retina. For blind patients suffering from photoreceptor degeneration, e.g. caused by diseases like Retinitis Pigmentosa, neuroprosthetic devices electrically stimulating remaining retinal neurons can restore basic visual abilities^77, 78^. The spatial and temporal resolution of such retinal implants is however still very limited^93^ with the highest reported restored visual acuity of 20/546^94^. While many experimental studies have explored different strategies of stimulation^95–98^, most of them are restricted to very specific stimulus types such as current or voltages pulses. As a consequence, retinal implants are not able to specifically target cell types such as the independent stimulation of the ON and OFF pathways of the retina^79, 99, 100^. To facilitate a systematic stimulus optimization in silico, we developed a simulation framework for electrical stimulation of the retina. While the idea to simulate responses of BCs to electrical stimuli is not new, previous studies restricted their models to point / ball source electrodes^6, 101^, simplified BCs to passive cables^102^ or used simplified BC models that only express L- or T-type channels^103^. Our framework combines the simulation of micro-electrode arrays used in neuroprosthetic devices^77, 78^ with detailed models of an OFF- and ON-BC. This allowed us to identify stimulus waveforms that can selectively target either OFF- or ON-BCs, which could help to better encode visual scenes into electrical signals of retinal implants. Likely, the different density for some ion channels contributed to the differential response of the two BC types. In addition, differences in morphology such as the axon length may have contributed as well. Ideally, in silicio optimized stimulation strategies would be first verified in ex vivo experiments before they could be implemented in neuroprosthetic devices to help to improve the quality of visual impressions.

To be able to simulate the electrical stimulation of the retina, we first inferred the conductivity and relative permittivity of the rd10 retina based on recorded currents evoked by sinusoidal stimulation voltages. While the estimated conductivity (*σ*_retina_ = 0.0764 S*/*m) is consistent with the value (*σ*_retina_ = 0.077 S*/*m) reported in ^80^, also smaller (0.025 S*/*m^104^) and larger (≈0.75 S*/*m^105^) conductivities have been reported for the retina. These differences may be due to different ways in tissue handling and preparation. Comparing the estimated values of the relative permittivity (*ε*_retina_ = 1.14×10^7^) to the literature is more difficult, and most simulation studies choose to ignore its effects. The relative permittivity of retinal tissue has been reported for very high frequencies (10 MHz to 10 GHz), but the strong frequency dependence makes a direct comparison to frequencies several orders of magnitude smaller (e.g. 40 Hz) not meaningful. Additionally, data from gray matter suggest a relative permittivity of 1.5×10^7^ at 40 Hz very close to our estimate^81^. This opens the question weather the common assumption to ignore the effects of the relative permittivity in other simulations^6, 102, 103^ is valid.

In summary, mechanistic models in neuroscience such as biophysically realistic multicompartment models have long been regarded as requiring many manual parameter choices or highly specific experiments to constrain them. We showed here how relatively standard, high-throughput imaging data in combination with likelihood-free inference techniques can be used to perform Bayesian inference on such models, allowing unprecedented possibilities for efficient optimization and analysis of such models. Importantly, this allow us to understand which parameters in such models are well constrained, and which are not, and determine which parameter combinations lead to similar simulation outcomes^17, 106^ — questions that have hindered progress in the field for years.

## Acknowledgements

We thank Yves Bernaerts for preliminary work on electrical stimulation of biophysical models, Timm Schubert for support with animal protocols and Gordon Eske and Adam vor Daranyi for technical support. Additionally, we thank Jakob Macke, Philipp Hennig and Lara Hoefling for discussion. This work was funded by the Federal Ministry of Education and Research (BMBF, FKZ 01GQ1601 and 01IS18052 to PB), the German Science Foundation through the Excellence Cluster 2064 “Machine Learning - New Perspectives for Science” (390727645), a Heisenberg Professorship (BE5601/4-1 to PB), the Baden-Württemberg Stiftung (NEU013 to PB and GZ), and the National Institutes of Health (NIH) (EY022070 and EY023766 to RGS).

## Author contributions

PB, TE, RS and GZ designed the study; JO, CB and RS performed modeling; JO and CS optimized ML algorithms; JO analyzed models and data; KF and TH performed experiments; TE and GZ supervised experiments; PB, GZ and CB supervised the project; JO and PB wrote the paper with input from all authors.

## Appendix

**Table 4.** Cone model parameter priors.

| Parameter | Unit | $a_i$ | $\mu_i$ | $b_i$ | Remarks | References |
| --- | --- | --- | --- | --- | --- | --- |
| $C_m$ | $\mu\text{F cm}^{-2}$ | 0.9 | 1 | 1.1 | | (22) |
| $R_m$ | $\text{k}\Omega \text{ cm}^2$ | 1 | 5 | 40 | | (22) |
| $V_r$ | mV | -80 | -65 | -50 | | |
| $\text{Ca}_L$ | $\text{mS cm}^{-2}$ | 0.1 | 2 | 10 | | (32,33,43) |
| $\text{K}_V$ | $\text{mS cm}^{-2}$ | 0 | 0.1 | 3 | | (36) |
| $\text{HCN @ } S$ | $\text{mS cm}^{-2}$ | 0.1 | 3 | 10 | | (29,35,36) |
| $\text{HCN @ } A$ | $\text{mS cm}^{-2}$ | 0.1 | 1 | 10 | | (29,35,36) |
| $\text{HCN @ } AT$ | $\text{mS cm}^{-2}$ | 0 | 0.1 | 10 | | (29,35,36) |
| $\text{Cl}_{\text{Ca}}$ | $\text{mS cm}^{-2}$ | 0 | 0.1 | 10 | | (38,39) |
| $\text{Ca}_P$ | $\mu\text{S cm}^{-2}$ | 0.1 | 10 | 100 | | (34) |
| $\tau_\alpha(\text{Ca}_L)$ | | 0.75 | 1 | 1.5 | | |
| $\tau_\alpha(\text{K}_V)$ | | 0.1 | 1 | 10 | | |
| $\Delta V_\alpha(\text{Ca}_L)$ | mV | -5 | 0 | 5 | | |
| $\Delta V_\alpha(\text{K}_V)$ | mV | -10 | 0 | 10 | | |
| $\text{Ca}_P\text{K}$ | $\mu\text{M}$ | 0.01 | 5 | 100 | | |
| RRP | vesicles | 10 | 20 | 30 |  | 58,59 |
| $g_l$ | | 0.3 | 1 | 3 | | |

**Table 5.**
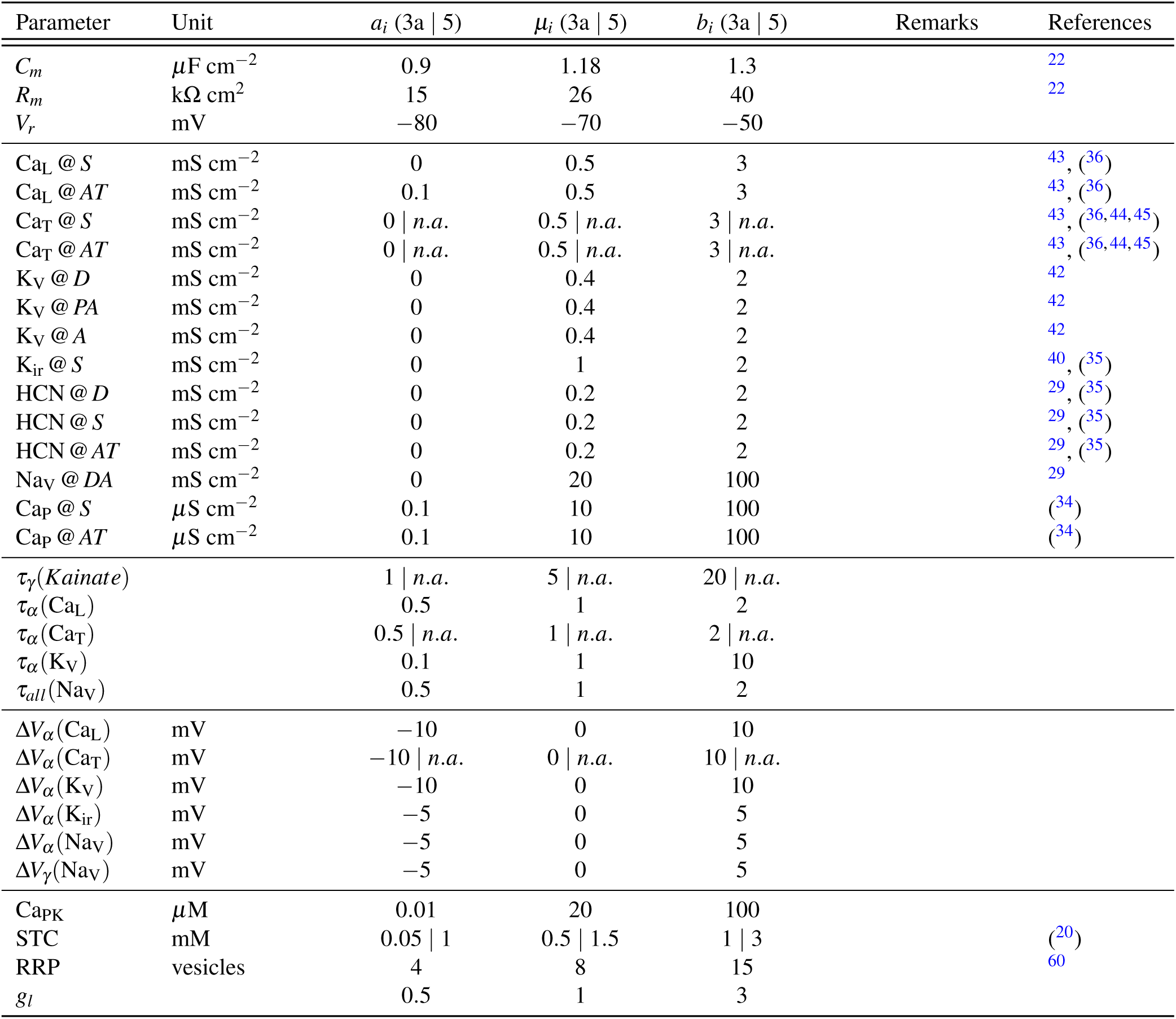
BC model parameter priors.

| Parameter | Unit | $a_i$ (3a \| 5) | $\mu_i$ (3a \| 5) | $b_i$ (3a \| 5) | Remarks | References |
| --- | --- | --- | --- | --- | --- | --- |
| $C_m$ | $\mu\text{F cm}^{-2}$ | 0.9 | 1.18 | 1.3 | | 22 |
| $R_m$ | $\text{k}\Omega \text{ cm}^2$ | 15 | 26 | 40 | | 22 |
| $V_r$ | mV | -80 | -70 | -50 | | |
| $\text{Ca}_L @ S$ | $\text{mS cm}^{-2}$ | 0 | 0.5 | 3 | | 43, (36) |
| $\text{Ca}_L @ AT$ | $\text{mS cm}^{-2}$ | 0.1 | 0.5 | 3 | | 43, (36) |
| $\text{Ca}_T @ S$ | $\text{mS cm}^{-2}$ | 0 \| *n.a.* | 0.5 \| *n.a.* | 3 \| *n.a.* | | 43, (36,44,45) |
| $\text{Ca}_T @ AT$ | $\text{mS cm}^{-2}$ | 0 \| *n.a.* | 0.5 \| *n.a.* | 3 \| *n.a.* | | 43, (36,44,45) |
| $\text{K}_V @ D$ | $\text{mS cm}^{-2}$ | 0 | 0.4 | 2 | | 42 |
| $\text{K}_V @ PA$ | $\text{mS cm}^{-2}$ | 0 | 0.4 | 2 | | 42 |
| $\text{K}_V @ A$ | $\text{mS cm}^{-2}$ | 0 | 0.4 | 2 | | 42 |
| $\text{K}_{ir} @ S$ | $\text{mS cm}^{-2}$ | 0 | 1 | 2 | | 40, (35) |
| $\text{HCN} @ D$ | $\text{mS cm}^{-2}$ | 0 | 0.2 | 2 | | 29, (35) |
| $\text{HCN} @ S$ | $\text{mS cm}^{-2}$ | 0 | 0.2 | 2 | | 29, (35) |
| $\text{HCN} @ AT$ | $\text{mS cm}^{-2}$ | 0 | 0.2 | 2 | | 29, (35) |
| $\text{Na}_V @ DA$ | $\text{mS cm}^{-2}$ | 0 | 20 | 100 | | 29 |
| $\text{Ca}_P @ S$ | $\mu\text{S cm}^{-2}$ | 0.1 | 10 | 100 | | (34) |
| $\text{Ca}_P @ AT$ | $\mu\text{S cm}^{-2}$ | 0.1 | 10 | 100 | | (34) |
| $\tau_\gamma(\text{Kainate})$ | | 1 \| *n.a.* | 5 \| *n.a.* | 20 \| *n.a.* | | |
| $\tau_\alpha(\text{Ca}_L)$ | | 0.5 | 1 | 2 | | |
| $\tau_\alpha(\text{Ca}_T)$ | | 0.5 \| *n.a.* | 1 \| *n.a.* | 2 \| *n.a.* | | |
| $\tau_\alpha(\text{K}_V)$ | | 0.1 | 1 | 10 | | |
| $\tau_{all}(\text{Na}_V)$ | | 0.5 | 1 | 2 | | |
| $\Delta V_\alpha(\text{Ca}_L)$ | mV | -10 | 0 | 10 | | |
| $\Delta V_\alpha(\text{Ca}_T)$ | mV | -10 \| *n.a.* | 0 \| *n.a.* | 10 \| *n.a.* | | |
| $\Delta V_\alpha(\text{K}_V)$ | mV | -10 | 0 | 10 | | |
| $\Delta V_\alpha(\text{K}_{ir})$ | mV | -5 | 0 | 5 | | |
| $\Delta V_\alpha(\text{Na}_V)$ | mV | -5 | 0 | 5 | | |
| $\Delta V_\gamma(\text{Na}_V)$ | mV | -5 | 0 | 5 | | |
| $\text{Ca}_{PK}$ | $\mu\text{M}$ | 0.01 | 20 | 100 | | |
| STC | mM | 0.05 \| 1 | 0.5 \| 1.5 | 1 \| 3 |  | (20) |
| RRP | vesicles | 4 | 8 | 15 |  | 60 |
| $g_l$ | | 0.5 | 1 | 3 | | |

**Table 6.** Other NeuronC parameters.

| Parameter | Unit | Value | Remarks |
| --- | --- | --- | --- |
| vna | mV | 65 | Reversal potential sodium |
| vk | mV | -89 | Reversal potential potassium |
| vcl | mV | -70 | Reversal potential chloride |
| dnao | mmol | 151.5 | Extracellular sodium concentration |
| dko | mmol | 2.5 | Extracellular potassium concentration |
| dclo | mmol | 133.5 | Extracellular chloride concentration |
| dcao | mmol | 2 | Extracellular calcium concentration |
| dicafrac |  | 1 | Fraction of calcium pump current that is added to total current |
| use_ghki |  | 1 | Use Goldman-Hodgkin-Katz equation for ion channels |
| cone_timec |  | 0.2 | Time constant of cone phototransduction |
| cone_loopg |  | 0.0 | Gain of calcium feedback loop in cones |
| cone_maxcond | nS | 0.2 | Maximum conductance of of outer segment |
| timinc | μs | 1 \| 5 \| 10 | Simulation time step (thresholds \| BC posterior samples \| otherwise) |
| ploti | ms | 0.01 \| 0.1 \| 2 | Recording time step (thresholds \| stimulus optimization \| otherwise) |
| stiminc | ms | 0.01 \| 0.1 | Synaptic time step (thresholds \| otherwise) |
| srtimestep | ms | 0.1 | Stimulus update time step |
| predur | s | 6 \| 3 | Simulation time to reach equilibrium potential (OFF \| ON) |

